# Single-nuclear transcriptomics of lymphedema-associated adipose reveals a pro-lymphangiogenic stromal cell population

**DOI:** 10.1101/2025.02.18.638907

**Authors:** Gregory P. Westcott, Margo P. Emont, Anton Gulko, Zhibo Zhou, Charissa Kim, Gopal Varma, Leo L. Tsai, Eliza O’Donnell, Zinger Yang Loureiro, Wenle Liang, Christopher Jacobs, Linus T. Tsai, Timothy P. Padera, Dhruv Singhal, Evan D. Rosen

## Abstract

Chronic lymphedema is a progressive, disfiguring disease that results from dysfunction of the lymphatic vasculature, causing distal accumulation of interstitial fluid, localized development of tissue edema, and expansion of subcutaneous adipose tissue (SAT). As the molecular mechanisms governing SAT remodeling in this disease are unclear, we performed single-nucleus RNA sequencing on paired control and affected SAT biopsies from patients with unilateral lymphedema. Lymphedema samples were characterized by expansion of SAA^+^ adipocytes, pro-adipogenic stem cells, and proliferation of lymphatic capillaries. A *GRIA1*^+^ lymphedema-enriched stromal cell population expressing *VEGFC*, *ADAMTS3*, and *CCBE1* was identified, suggesting an enhanced axis of communication between adipose stem and progenitor cells (ASPCs) and lymphatic endothelial cells. Furthermore, lymphedema ASPC-conditioned media promoted lymphatic endothelial tube elongation and proliferation *in vitro*. These findings indicate a critical role for ASPCs in regulating adipocyte differentiation and lymphatic vascular remodeling in lymphedema, and provide a valuable resource for better understanding this disease.

## Background

Lymphedema is a progressive condition that results from impaired lymphatic drainage of the subcutaneous interstitial compartment, leading to fluid accumulation localized to the distal anatomic region^1^. With chronic lymphatic insufficiency, adipose differentiation takes place in the affected distribution^2^, rendering the swelling irreversible, negatively impacting quality of life^3^ and predisposing to other skin complications such as recurrent infection^4^. In developed geographic regions, lymphedema is most often a complication of surgical lymphadenectomy or radiation for cancer treatment, affecting at least 20% of women undergoing treatment for breast cancer^5^. Worldwide, filariasis infection is the most prevalent etiology^6^. Obesity is also an increasingly common risk factor for lymphedema^7^, particularly when the body mass index surpasses 40 kg/m^2^. Despite a worldwide prevalence estimated at 140-250 million^6^, there are limited non-surgical options to prevent, slow, or reverse lymphedema. Adjunctive medical approaches, such as VEGF-C-based adenoviral therapy^8^ or administration of the leukotriene B4 inhibitor ketoprofen^9^, have failed to show improvement in lymphedema volume or lymphoscintigraphy parameters, highlighting the urgent need for novel therapeutic targets to address the primary burden of disease.

SAT expansion that occurs in chronic lymphedema confers much of the symptomatic disease burden and reflects the complex relationship between the lymphatic system and adipocyte differentiation and function^10,11^. In order to better characterize the complex interplay of diverse cell types, including adipocytes, and identify disease-specific cell states and cellular interactions, we performed single-nucleus RNA sequencing (sNuc-seq) on lymphedema-associated SAT, an approach we and others have previously used to construct a detailed atlas of human and mouse adipose tissue^12–15^. We also took advantage of the unilateral nature of localized limb lymphedema to obtain both affected and control samples from corresponding sites of the same subject, eliminating the confounding effects conferred by comparing two separate groups of individuals, and significantly improving our power to detect subtle differences between groups. Furthermore, sNuc-seq allows for the sequencing of mature adipocytes and the improved recovery of endothelial populations when compared to whole-cell sequencing technologies^12^. Our sequencing data and subsequent experimental studies indicate a role for adipose stem and progenitor cells (ASPCs) in coordinating both adipocyte differentiation as well as lymphatic capillary remodeling.

## Results

### Single-nucleus RNA sequencing atlas of lymphedema-associated SAT

Subjects with unilateral, chronic, fat-dominant lymphedema undergoing debulking surgery were recruited pre-operatively from the Beth Israel Deaconess Medical Center Department of Plastic and Reconstructive Surgery. Paired SAT biopsies were obtained from corresponding sites on the lymphedematous and unaffected (control) limbs from seven subjects for sNuc-seq, and lipoaspirate was collected from lymphedema and control subjects for ASPC isolation (**Figure 1a**). Samples were obtained from anatomical sites where lymphedema-associated adipose deposition is typically most substantial, specifically the posterior aspect of the calf approximately 5 centimeters distal to the popliteal fossa, or the posterior aspect of the upper arm approximately 5 centimeters proximal to the olecranon. See **Supplementary Table 1** for additional subject details. Only one subject had primary lymphedema, and one subject was male, limiting our ability to compare samples by etiology or sex. Magnetic resonance imaging (MRI) features were determined via intensity-based threshold segmentation at the approximate location of each biopsy, and revealed expected increases in total subcutaneous area, subcutaneous fat, and interstitial fluid cross-sectional area in lymphedema (**Figure 1b-c** and **Supplementary Table 2)**.

**Figure 1.**
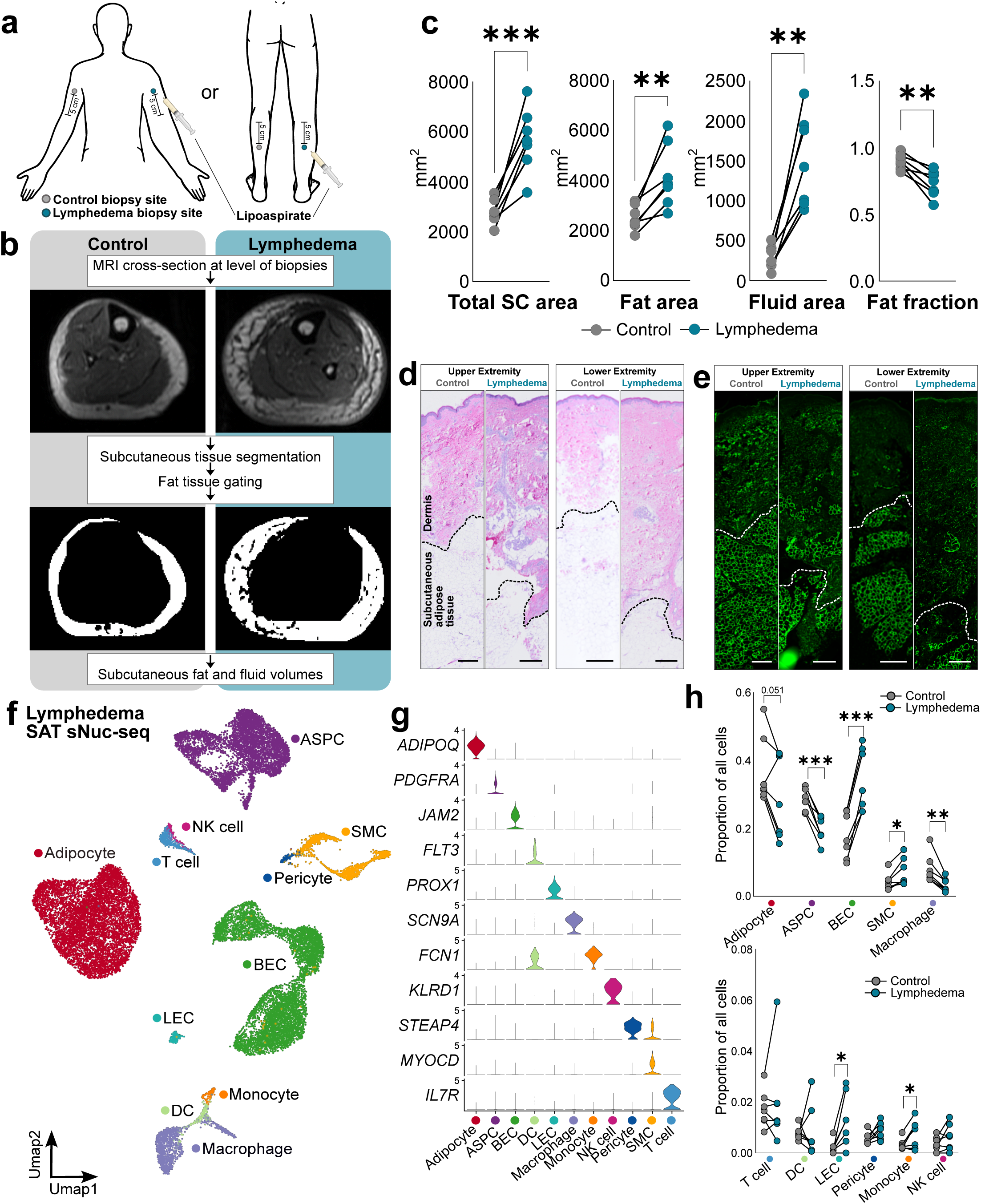
Single-nucleus RNA sequencing atlas of lymphedema-associated subcutaneous adipose tissue. *a,* Tissue collection sites; biopsies were taken approximately 5 cm proximal to epicondyle of the humerus or 5 cm distal to the popliteal fossa; *b,* MRI analysis workflow and example images; *c*, Quantification of MRI parameters, n=7 pairs **(Supplementary Table 2)**; *d*, Hematoxylin and eosin staining of control and lymphedema biopsies; *e*, Perilipin 1 immunofluorescence of control and lymphedema biopsies; *f*, UMAP of lymphedema sNuc-seq atlas, including 23,777 cells; *g*, Log-transformed expression of marker genes for cell clusters identified by sNuc-seq; *h*, Proportions of cell types per individual sample, n=7 pairs of samples (**Supplementary Table 3**). ASPC, adipose stem and progenitor cell; BEC, blood endothelial cell; DC, dendritic cell; LEC, lymphatic endothelial cell; NK: natural killer; SC: subcutaneous; SMC: smooth muscle cell; Statistical analysis performed using paired two-tailed Student’s t-test: *, p<0.05; **, p<0.01; ***, p<0.001

*En bloc* SAT biopsies were processed for sNuc-seq and histology. **Figure 1d-e** illustrates the anatomic relationship of the SAT, which was used for all analyses, to the dermis, which becomes thickened in chronic lymphedema^16^. A total of 23,777 cells (10,785 control and 12,992 lymphedema) were identified in the sNuc-seq dataset, which includes all expected major adipose tissue cell types (**Figure 1f-g, Extended Data Figure 1, and Supplementary Table 3**). Notably, there was a relative increase in vascular cells in lymphedema samples, reflected by an increase in blood and lymphatic endothelial cells as well as smooth muscle cells, while ASPCs, adipocytes, and macrophages decreased in proportion (**Figure 1h**).

The decrease in the proportion of macrophages was unexpected given prior evidence that macrophages accumulate within the dermis in lymphedema^17^. Macrophages were subset into Mac1 and Mac2 subpopulations (**Extended Data Figure 2a-b**), and the proportional decrease in macrophages was mainly driven by a reduction in the Mac1 subpopulation, which is characterized by expression of *LYVE1* and M2-like markers *CD163* and *MRC1* (CD206), while Mac2, which expresses *TREM2* and lipid handling genes such as *LPL* and *FABP4,* did not change in proportion (**Extended Data Figure 2c-d**). Though LYVE1+ macrophages promote angiogenesis^18^, they also recede in the setting of acute injury and inflammation with subsequent reconstitution during tissue healing^19^, and their dynamics may therefore be altered in ongoing tissue injury due to chronic lymphedema. Furthermore, macrophages did not significantly express *VEGFA* or *VEGFC* (**Extended Data Figure 2c**), indicating that, unlike has been reported in mouse models of lymphedema^17^, macrophages are not likely to be the key factor driving the angiogenesis observed in our samples.

### Lymphedema is characterized by expansion of an SAA^+^ adipocyte population and smaller adipocyte size

Adipocytes were assigned to six subpopulations and labeled according to mapping with previously-defined subcutaneous human adipocyte subclusters^12^ (**Figure 2a-c and Extended Data Figure 3a-b**). Ad^TNS1^ expresses tensin 1, which is a marker of the previously described Adipo^Lep^ population^14^, though leptin expression is similar to that of Ad^SAA^ (**Extended Data Figure 3c**). Ad^ADGRB3^ is a small population that expresses adhesion G protein-coupled receptor B3 (also known as brain angiogenesis inhibitor-3, *BAI3*), which was recently noted to have a role in adaptive thermogenesis^20^. Ad^SAT1^ expresses spermidine/spermine N1-acetyltransferase 1, which performs polyamine catabolism required for adipocyte thermogenesis; knockout of *Sat1* in adipocytes results in obesity^21^. Ad^ITGBL1^ expresses integrin-β–like 1, an integrin inhibitor that promotes chondrogenesis, as well as collagen gene *COL6A2*. Ad^Basal^ lacked specific marker genes to distinguish it from other adipocyte populations.

**Figure 2.**
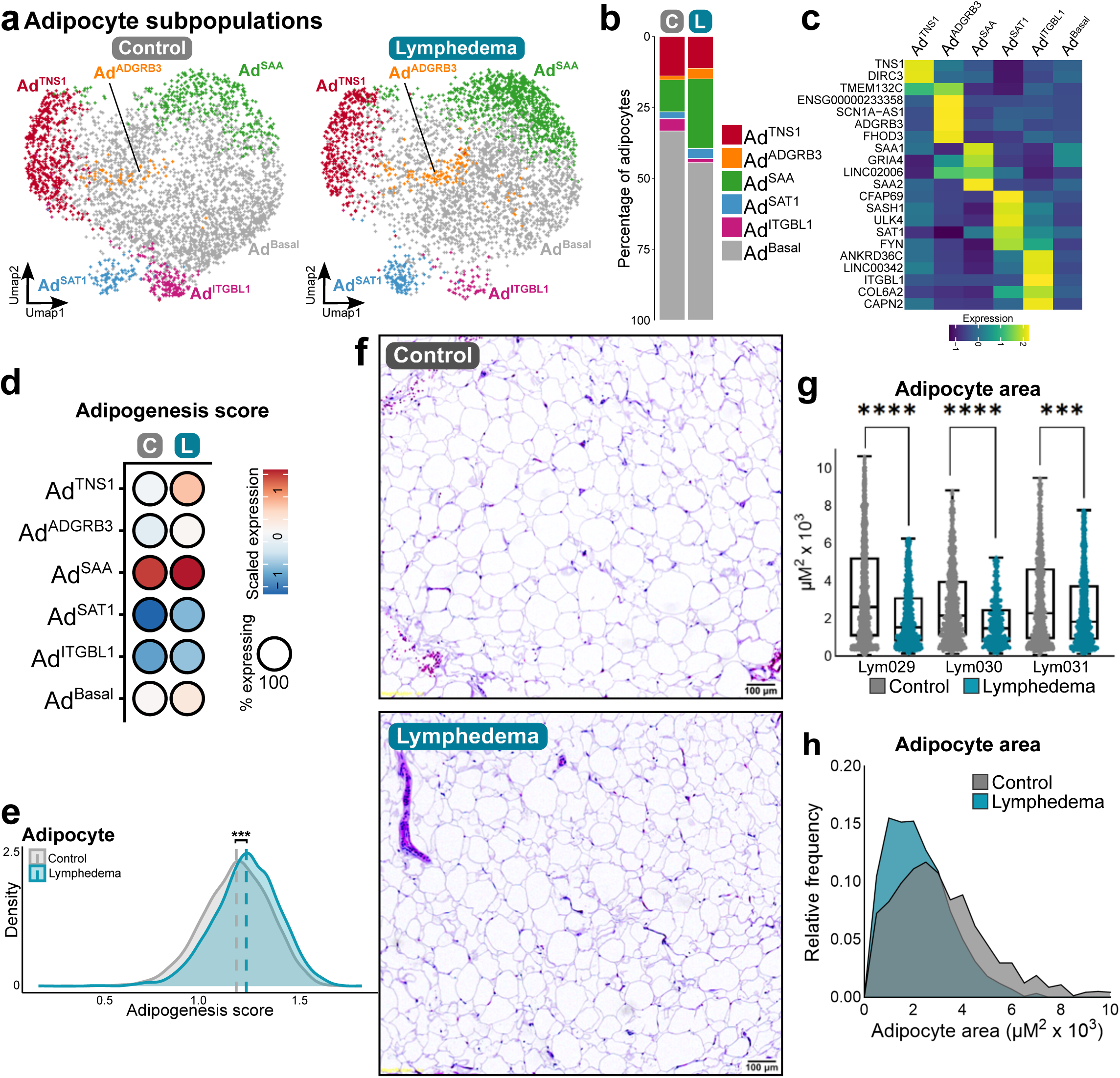
Lymphedema is characterized by expansion of *SAA^+^* adipocytes and smaller adipocyte size. Adipocyte (*a-c*) subpopulations, relative proportions, and scaled expression of marker genes; *d*, Adipogenesis scores disaggregated by adipocyte subpopulation and disease state; *e*, Adipogenesis score density plot across all adipocytes; *f*, Hematoxylin and eosin staining of representative SAT samples; *g*, Adipocyte size quantification across paired samples; *h*, Distribution of adipocyte size; C, control; L, lymphedema. Statistical analysis performed using Wilcoxon rank-sum test (e) or paired two-tailed Student’s t-test (g); ***, p<0.001; ****, p<0.0001

Ad^SAA^ was of particular interest, as this subpopulation is significantly greater as a proportion of adipocytes in lymphedema (23.5%) compared to control (11.1%) (*p* = 0.013) (**Figure 2b** and **Extended Data Figure 3d**). Ad^SAA^ expresses acute phase serum amyloid A isoforms *SAA1* and *SAA2* (**Extended Data Figure 3e)**, which are apolipoproteins that are upregulated with increasing adipocyte size^22,23^ in obesity^24^ and during adipogenesis^25^. These serum amyloid A isoforms may play a role in cholesterol efflux to macrophages from hypertrophied adipocytes^22^. An adipogenesis gene signature (Hallmark Adipogenesis)^26^ was applied to adipocyte and ASPC populations to assess the degree of differentiation^27^, identifying Ad^SAA^ as the most differentiated adipocyte subpopulation (**Figure 2d**). Furthermore, there was an increased overall adipocyte adipogenesis score in lymphedema across all adipocytes in lymphedema (**Figure 2e**) as well as when assessed on a per-individual basis (**Extended Data Figure 3f**). Only one adipocyte sample did not increase in adipogenesis score, which happened to be the sole male sample collected.

Adipocyte size was measured within the subcutaneous area (indicated in Figure 1d-e) of paired biopsy samples from three subjects on H&E images, and confirmed with a fourth subject using BODIPY staining of whole mounted SAT samples. In all subjects, average adipocyte size was decreased in lymphedema (**Figure 2f-h** and **Extended Data Figure 3h**). The literature describing adipocyte size in lymphedema is mixed, with some studies finding larger adipocytes in lymphedema^28,29^, while others found no difference^30^ or a lower proportion of hypertrophied adipocytes in lymphedema SAT^31^, similar to our results. The presence of smaller adipocytes suggests the increase in fat volume may be driven by substantial adipocyte hyperplasia and ongoing progenitor differentiation into newly-formed adipocytes.

### Lymphedema-associated ASPCs increase adipogenic and proliferative gene expression

In order to further investigate the response of adipocyte progenitors to lymphedema, ASPCs were subclustered into six populations (**Figure 3a-c** and **Extended Data Figure 3i-j**). ASPC^NAMPT^ is characterized by expression of the NAD+ synthesis enzyme nicotinamide phosphoribosyltransferase, which has been implicated in adipose tissue plasticity and insulin resistance^32–34^, as well as by expression of a number of collagen genes. ASPC^CD81^ expresses CD81, indicating it could represent a beige adipocyte progenitor population^35^. ASPC^CD55^ represents early progenitor cells expressing dipeptidyl peptidase 4 (*DPP4*) and *CD55*^36^. ASPC^PTCH2^, which increased modestly from 7.9% to 11.9% in lymphedema, expresses hedgehog receptor patched 2 and shares transcriptional features with anti-adipogenic Aregs^37^. Notably, lymphedema is characterized by the dramatic increase in ASPC^GRIA1^; ASPC^GRIA1^ is enriched for expression of the essential lymphangiogenic ligand *VEGFC* and the surface receptor glutamate receptor 1 (*GRIA1*).Like adipocytes, ASPCs expressed a significantly higher adipogenesis signature in lymphedema (**Figure 3d-e** and **Extended Data Figure 3f-g**), consistent with a predisposition toward adipocyte differentiation. Most subpopulations of ASPCs also exhibited an increased stem cell proliferation signature (**Figure 3f**).

**Figure 3.**
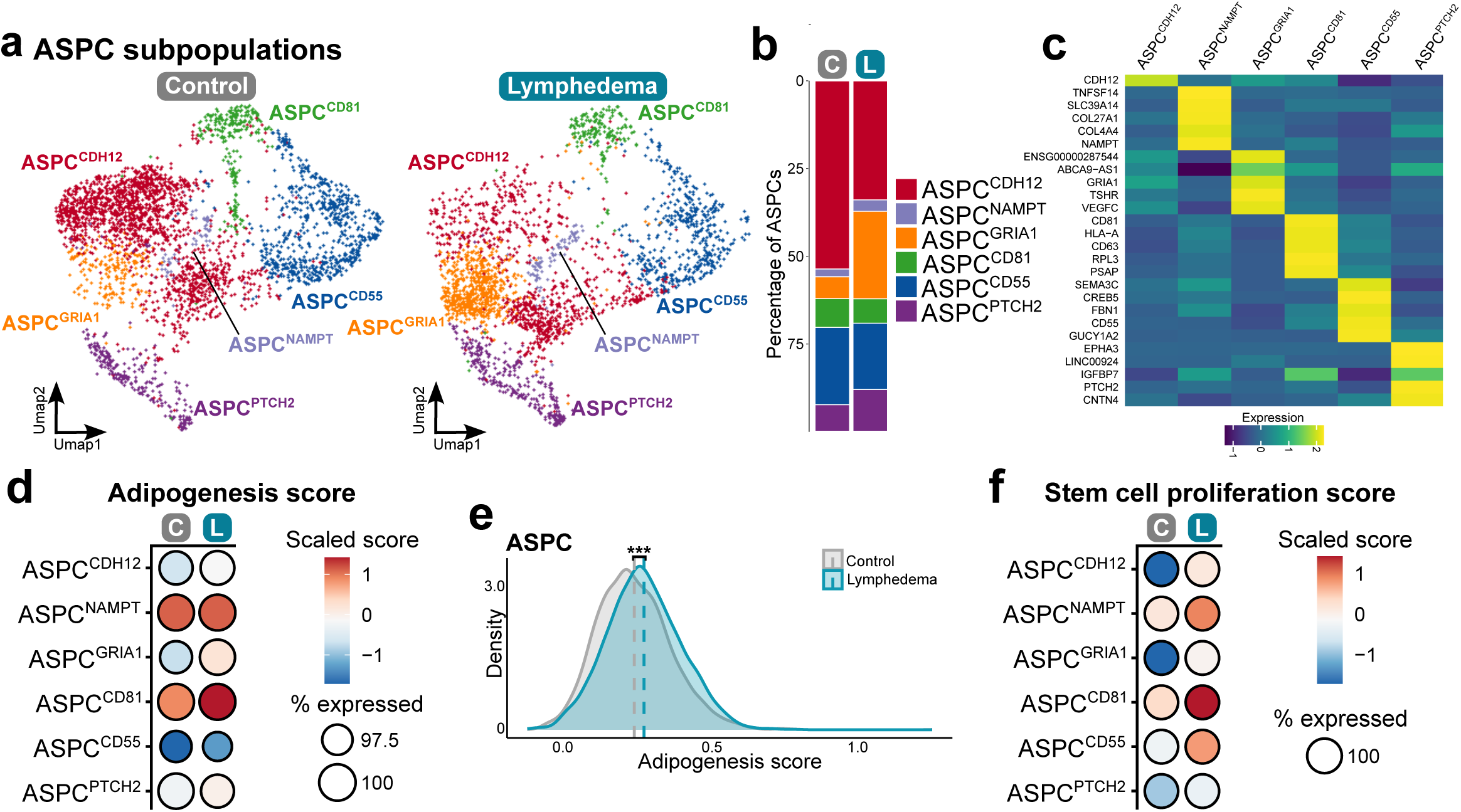
ASPCs exhibit adipogenic and proliferative gene expression in lymphedema. ASPC (*a-c*) subpopulations, relative proportions, and scaled expression of marker genes; *d*, Adipogenesis score by subpopulation and *e*, across all ASPCs; *f*, Stem cell proliferation score by ASPC subpopulation. C, control; L, lymphedema; Statistical analysis performed using Wilcoxon rank-sum test; ***, p<0.001

### ASPCs upregulate a subset of genes that correlate with disease characteristics and demonstrate augmented WNT, IGF, and BMP signaling in lymphedema

To further characterize the ASPC response to lymphedema, pseudobulk analysis was performed. In order to mitigate variability related to cell subsampling, pseudobulk was run 10,000 times, and differentially expressed genes (DEGs) that met significance in 90% of iterations were selected for further analysis (**Figure 4a**). Pearson correlation coefficients were determined between DEGs and MRI characteristics to identify genes associated with disease severity, and subsets of genes that significantly correlated with subcutaneous fat volume, fluid volume, or total volume were identified (**Figure 4b and Extended Data Figure 4**). ASPC subpopulations expressed and upregulated distinct patterns of these volume-associated genes; for example, ASPC^CDH12^ displayed upregulation of genes that were positively associated with both fluid and fat volume, while ASPC^NAMPT^ preferentially upregulated fluid-associated genes, and ASPC^GRIA1^ upregulated fat-associated genes (**Figure 4c**). ASPC^PTCH2^ did not appear to significantly express either fat- or fluid-associated genes. ASPC subpopulations therefore may play unique roles in responding to interstitial fluid accumulation and mediating the natural history of lymphedema.

**Figure 4.**
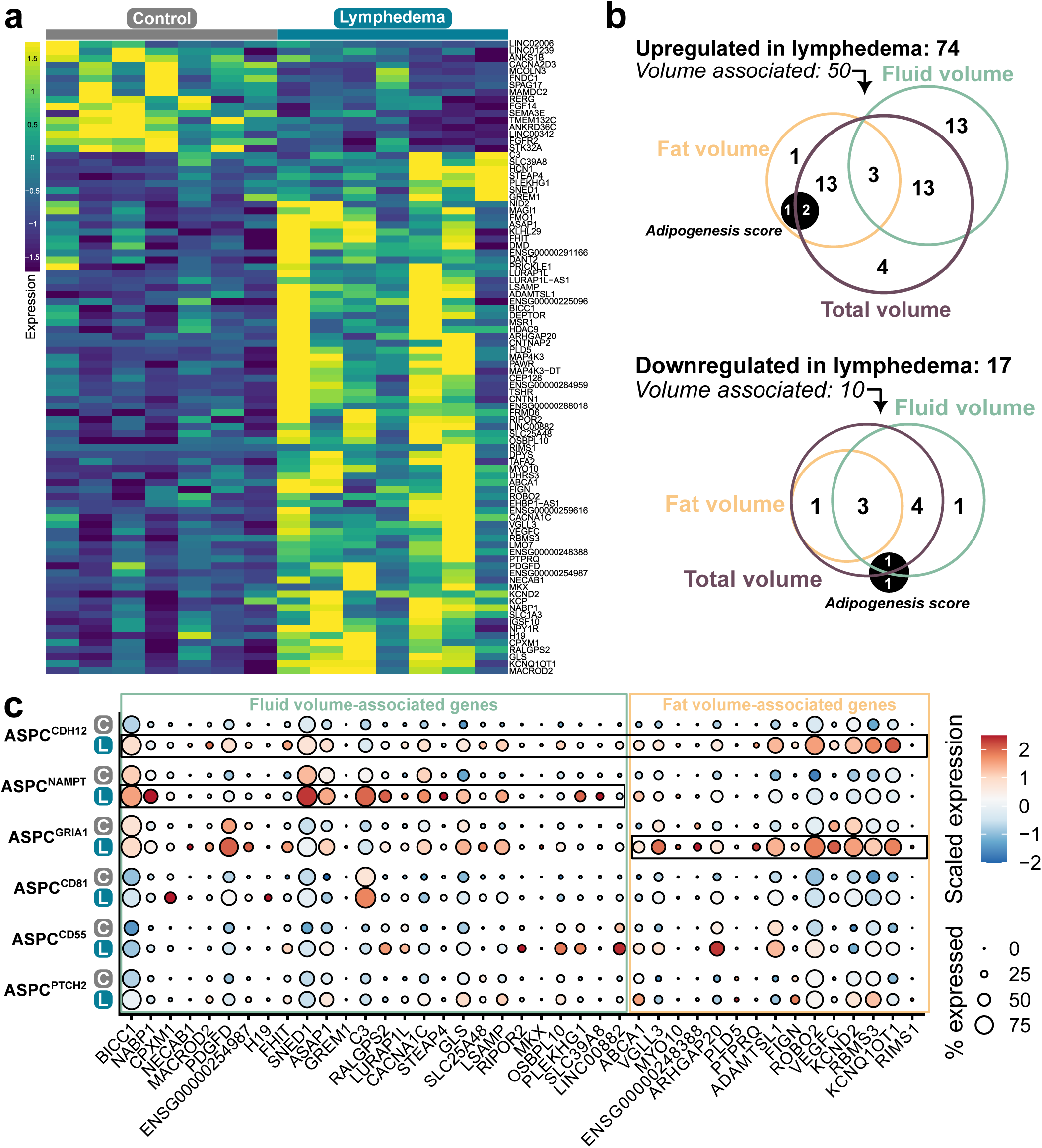
Lymphedema-associated ASPCs upregulate a subset of genes that correlate with disease characteristics. *a*, Heatmap of significant genes from ASPC pseudobulk analysis by sample; *b*, Categorization of pseudobulk genes that were significantly associated with either fat, fluid, or total subcutaneous area; *c*, Positively correlated volume-associated gene expression by ASPC subpopulation. C, control; L, lymphedema;

NicheNet^38^ was applied to the gene expression profiles of ASPC subpopulations to model intercellular communication and identify regulatory signaling pathways. The top induced signaling pathways in lymphedema included *WNT1*, *NTF4*, *IGF2*, *BMP2, and TGFB1* (**Figure 5a** and **Extended Data Figure 5a**). Applying signaling gene signatures on a per-cell basis with Vision, it became apparent that ASPC^CD55^, the DPP4+ stem cell population, activates signaling across multiple pathways, while more specialized populations are more selectively regulated (**Fig. 5b**). In lymphedema, WNT was upregulated across all subtypes, while other pathways showed subpopulation-specific expression patterns. Canonical Wnt signaling has been implicated in preserving ASPC proliferative capacity and inhibiting adipogenesis^39^, and was also upregulated in most ASPC subpopulations in lymphedema. The coexistence of both pro-adipogenic and proliferation signatures across ASPC subpopulations may reflect the demand to continuously replenish adipocyte progenitor cells in the face of chronically dysregulated adipocyte hyperplasia.

**Figure 5.**
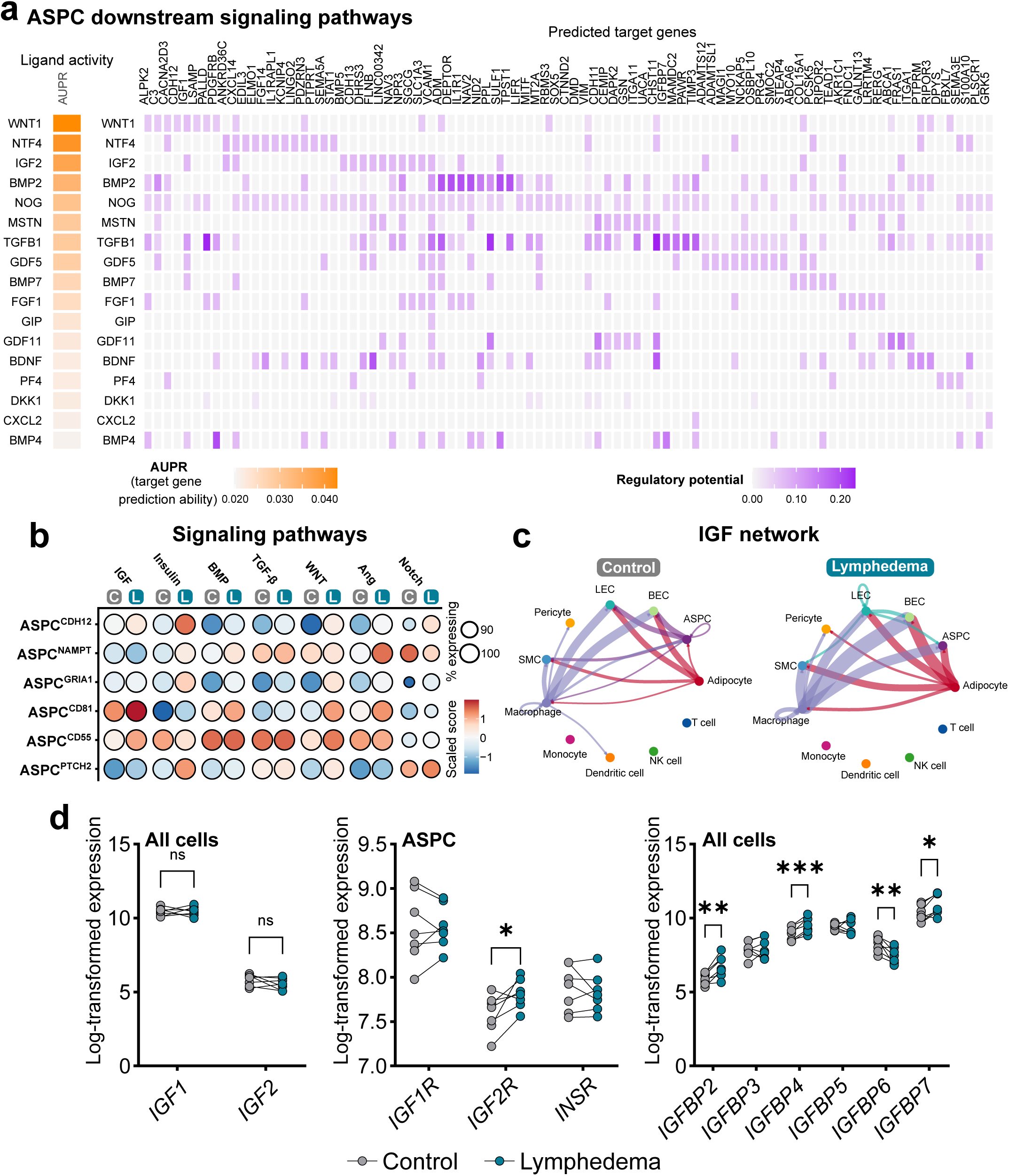
ASPCs exhibit altered WNT and IGF signaling in lymphedema. *a*, NicheNet (sender-agnostic) modeling of ASPC receptor-mediated signaling pathways increased in lymphedema; *b*, Selected signaling signature scores calculated by Vision, grouped by ASPC subpopulation; *c*, CellChat intercellular interaction modeling of the IGF signaling network between cell types; *d*, Pseudobulk expression of IGFs, IGF receptors, and IGFBPs. AUPR, area under the precision-recall curve. Statistical analysis performed using paired two-tailed Student’s t-test: ns, not significant; *, p<0.05; **, p<0.01; ***, p<0.001

IGF-2 protein concentration increases in lymphedema fluid^40^ and induces differentiation of subcutaneous preadipocytes^41^; the IGF pathway was most strongly upregulated in ASPC^CD81^ and ASPC^CD55^. CellChat^42^ ligand-receptor modeling predicted increased IGF signaling to ASPCs originating from macrophages, lymphatic endothelial cells (LECs), and adipocytes in lymphedema (**Figure 5c**). Though *IGF1* and *IGF2* are not upregulated on a whole-tissue level, *IGF2R* is upregulated in ASPCs, as are IGF binding proteins *IGFBP2*, *IGFBP4*, and *IGFBP7* across the entire tissue (**Figure 5d**). IGFBPs regulate adipocyte differentiation via both IGF-dependent and -independent mechanisms^43^, and downregulation of IGFBP5 and subsequent enhanced IGF signaling resulted in abnormal LEC remodeling in a mouse model of lymphedema^44^. Notably, IGFBP4 is essential for adipogenesis in mice^45^, while IGFBP7 promotes bovine^46^ and chicken^47^ preadipocyte differentiation.

BMP-2 also enhances adipogenesis^48^, and BMP signaling was upregulated in ASPC^NAMPT^ and ASPC^CD81^, with a predicted blood endothelial cell (BEC) source (**Extended Data Figure5b**). TGF-β1, which inhibits adipocyte differentiation but promotes collagen formation in models of lymphedema^49^, was induced in both ASPC^PTCH2^, which maps to anti-adipogenic Aregs, and ASPC^NAMPT^, which expresses *COL271A1* and *COL4A4* (**Extended Data Figure 5c**). Altogether, these data indicate that the ASPC response to lymphatic insufficiency is diverse and mediated by multiple distinct subpopulations.

### A unique GRIA1^+^ population of ASPCs expands dramatically in lymphedema and expresses critical lymphangiogenic ligands

As part of our effort to link specific cell populations in the adipose niche to the pathology of lymphedema, we asked which cell types express genes that have been shown to cause primary lymphedema when mutated. As expected, most such genes show high or even dominant expression in LECs. We were surprised to note, however, that two lymphedema genes, *CCBE1* and *ADAMTS3*, are almost exclusively expressed by ASPCs (**Figure 6a**). *CCBE1* and *ADAMTS3* encode components of the proteolytic cleavage machinery that activates VEGF-C to promote lymphangiogenesis^50–52^, and when mutated, cause Hennekam lymphangiectasia–lymphedema syndrome^53,54^. In lymphedema subjects, ASPC *VEGFC* expression is positively correlated with fat deposition (**Figure 6b**), and *VEGFC*, *ADAMTS3*, and *CCBE1* are co-expressed by the ASPC^GRIA1^ population (**Figure 6c-d**), which expands from 5.8% to 24.4% of total ASPCs (*p* < 0.001) (**Figure 6e**). Accordingly, CellChat predicted increased VEGF network communication from ASPCs to LECs in lymphedema (**Figure 6f**). ASPC^GRIA1^ also specifically expresses additional secreted ligands which have been demonstrated to have pro-lymphangiogenic activity, including *SVEP1*^55,56^ and *PDGFD*^57^ (**Extended Data Figure 5d**). In zebrafish, a population of fibroblasts co-express *vegfc, adamts3*, and *ccbe1* in proximity to lymphatic precursors during development^58^, suggesting that a stromal pro-lymphatic program is highly conserved and essential for lymphangiogenesis.

**Figure 6.**
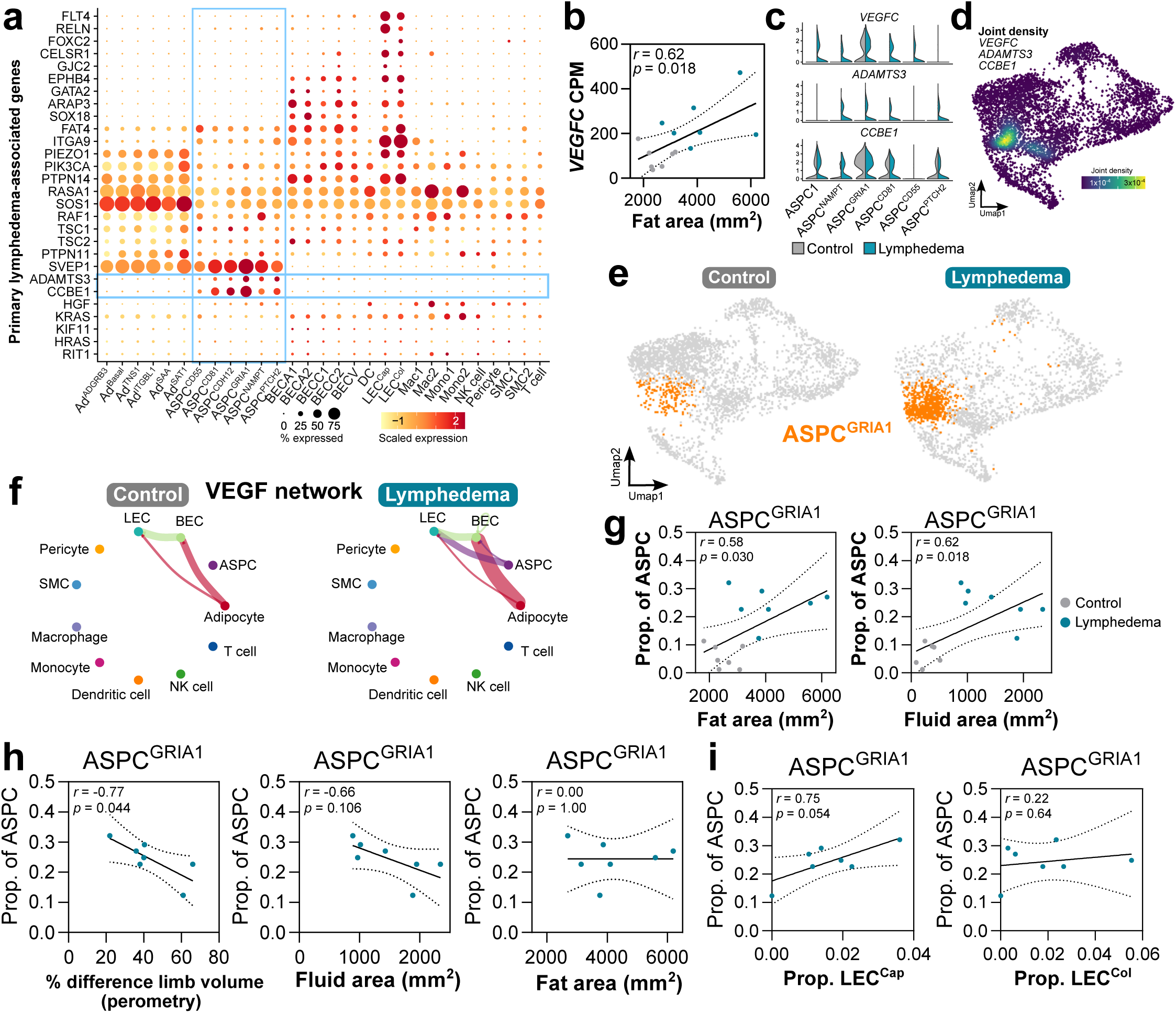
GRIA1^+^ ASPCs expand dramatically in lymphedema and express essential lymphangiogenic ligands. *a*, Selected genes associated with primary lymphedema syndromes plotted against cell subpopulations; *b*, *VEGFC* expression correlation with fat volume by MRI; *c*, Violin plots of lymphangiogenic gene expression within ASPC subpopulations; *d*, Joint density plot of VEGF-C related gene expression; *e*, UMAP plot of ASPC^GRIA1^ population in control and lymphedema; *f*, CellChat plot of predicted VEGF network ligand-receptor interactions; *g*, Correlation between all samples of ASPC^GRIA1^ proportion with MRI-derived fat and fluid areas; *h*, Correlation between lymphedema samples only of ASPC^GRIA1^ proportion with MRI-derived fat and fluid and *i*, proportions of LEC subpopulations.

As expected, the proportion of ASPC^GRIA1^ was positively associated with both fat and fluid areas by MRI (**Figure 6g**). Interestingly, within lymphedema limb samples, however, ASPC^GRIA1^ proportion was inversely correlated with severity of lymphedema as measured by perometry and trended toward a negative association with fluid area by MRI, but had no association with fat area (**Figure 6h**). ASPC^GRIA1^ proportion was also positively correlated with the proportion of LEC^Cap^, but not LEC^Col^ (**Figure 6i**).

Altogether, these findings indicate that ASPC^GRIA1^ proportion is associated with increased lymphatic density, decreased limb size, and possibly lower interstitial fluid volume.

### Lymphedema SAT exhibits active blood and lymphatic angiogenesis

BECs and LECs were both significantly increased in lymphedema (**Figure 1h**). Subclustering these populations revealed similar endothelial subpopulations to those previously described^12^ (**Figure 7a and Extended Data Figure 6a-c**), including lymphatic capillary (LEC^Cap^) and collecting duct (LEC^Col^) populations, which were delineated with recently described markers of skin lymphatics^59^ (**Figure 7b**). Of these subpopulations, only LEC^Cap^ was significantly enriched as a proportion of total endothelial cells in lymphedema (**Figure 7c** and **Extended Data Figure 6b**). Subpopulations BECV2, one of two *ACKR1*+ venous populations^60^, and LEC^Col^ demonstrated the highest angiogenesis scores calculated (**Figure 7d**). The overall angiogenesis score was higher in lymphedema for both BECs and LECs (**Figure 7e**).

**Figure 7.**
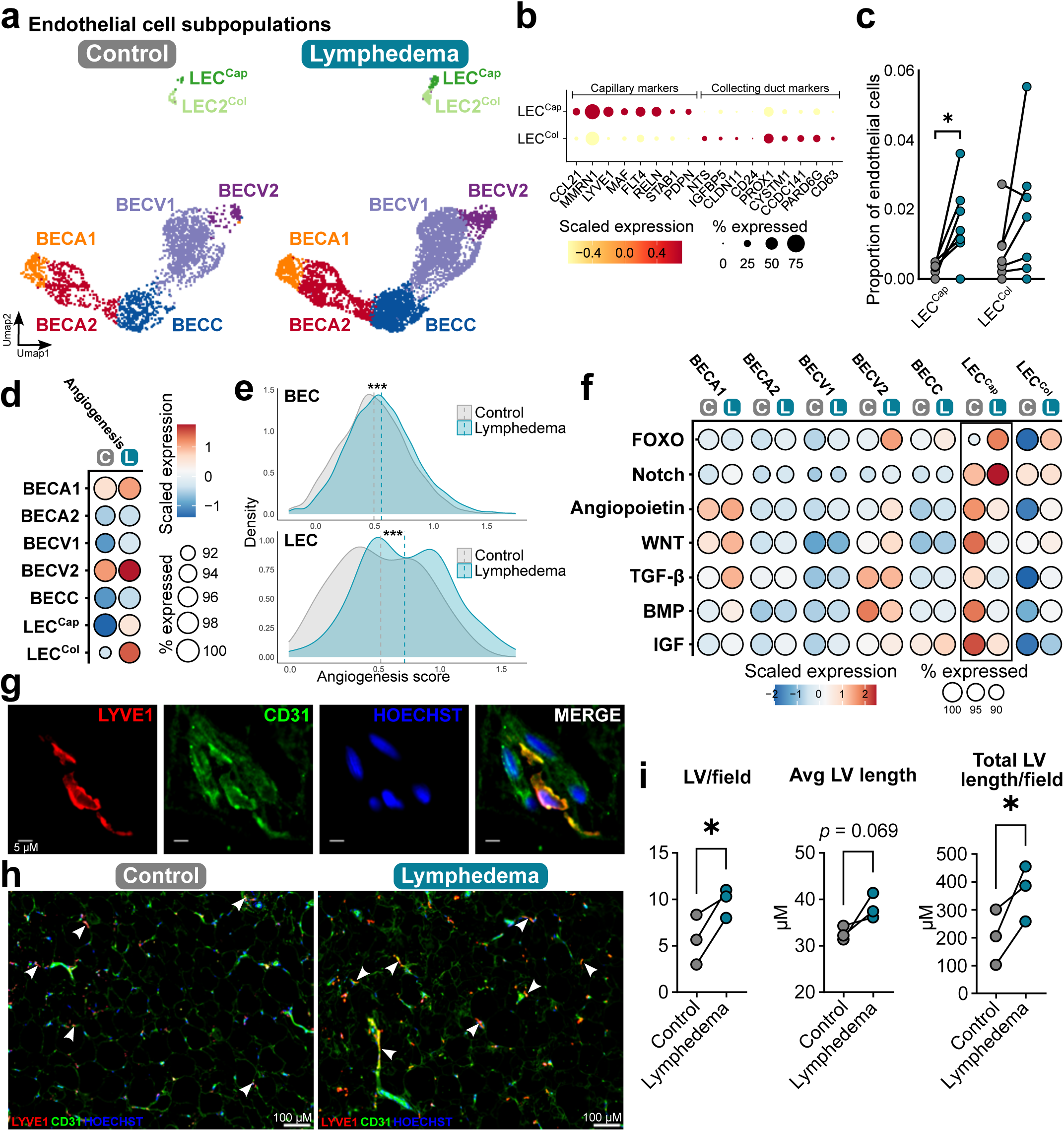
Lymphedema-associated SAT exhibits lymphangiogenesis. *a*, UMAP plots of endothelial populations split by disease status; *b*, Lymphatic capillary and collecting duct endothelial cell markers; *c*, LEC subpopulations as a proportion of total endothelial cells; *d*, Angiogenesis signature scores by subpopulation and *e*, across total BEC and LEC populations; *f*, Signaling pathway scores by endothelial subpopulation; *g*, Immunofluorescence of a representative lymphatic capillary and *h*, of a lower-power SAT field (arrow heads: LYVE1^+^CD31^+^ LVs); *i*, Quantification of SAT lymphatic vessel parameters by immunofluorescence of paired biopsies. LV, lymphatic vessels; C, control; L, lymphedema. Statistical analysis performed using Wilcoxon rank-sum test (e) or paired two-tailed Student’s t-test (c, i); *, p<0.05

Lymphedema-associated adipose tissue is characterized by hypoxia^61^, leading to alterations in expression of HIF-1α and HIF-2α (encoded by *EPAS1*). Our data reproduces previous reports that LEC HIF-1α expression is increased in lymphedema while HIF-2α may be mildly reduced (**Extended Data Figure 6d**), a pattern that has been shown to exacerbate lymphatic remodeling in lymphedema^62^ and may have implications for disease pathophysiology. HIF-1α also directly stimulates *VEGFA* expression^63^. Notably, the adipocyte population that expands in lymphedema, Ad^SAA^, highly expresses both *HIF1A* and *VEGFA* (**Extended Data Figure 6e**), presenting a possible mechanism contributing to the increase in blood endothelial cells.

A lymphedema-associated switch in signaling pathways was particularly evident in lymphatic capillaries, as FOXO and Notch were induced, and angiopoietin, Wnt, TGF-β, BMP, and IGF pathways were relatively downregulated (**Figure 7f**, box). FOXO1 is critical for developmental lymphangiogenesis^64^, while Notch1 activation has been shown to be essential in performing postnatal pathologic lymphangiogenesis^65^. TGF-β1^66^ and BMPs, such as BMP-9^67^, are anti-lymphangiogenic, while IGF^68^, Wnt^69^, and angiopoietin^70^ typically play pro-lymphangiogenic roles, suggesting a complex balance of lymphatic remodeling is at work in lymphedema, particularly considering lymphatic collectors demonstrate upregulation of these pathways.

SAT biopsy sections stained for CD31 revealed foci of increased blood vascularization in lymphedema and increased vascular density (**Extended Data Figure 6f-g**). Quantification was performed on vasculature present within SAT only, and does not include the dermis. Labeling for LYVE1 and CD31 co-expression also confirmed the presence of lymphatic vessels within SAT (**Figure 7g-h**). Quantification of paired biopsy sections demonstrated an increase in the number, average length, and total lymphatic length in lymphedema SAT (**Figure 7i**), consistent with our sequencing data as well as prior literature^37^. Whether functional lymphatic vessels are being formed, however, remains unclear.

### Lymphedema ASPC-secreted factors promote lymphatic endothelial tube elongation and LEC proliferation *ex vivo*

Given the presence of a lymphedema-selective ASPC subpopulation that expresses pro-lymphatic secreted ligands in the setting of increased lymphatic vascularization, we sought to determine if ASPCs isolated from lymphedema patients exert a cell-autonomous pro-lymphangiogenic effect. We performed a matrigel tube formation assay using primary human dermal lymphatic endothelial cells (HDLECs) treated with culture media conditioned by ASPCs isolated from either unaffected control or lymphedema SAT (**Figure 8a** and **Extended Data Figure 7a**). Treatment of HDLECs with lymphedema ASPC-conditioned media resulted in fewer, longer endothelial tubes (**Figure 8b**) as well as increased HDLEC proliferation (**Figure 8c**). Conditioned media treatment of human umbilical vein endothelial cells (HUVECs), however, had no impact on any tube formation parameters (**Extended Data Figure 7b**), suggesting a lymphatic-specific effect.

**Figure 8.**
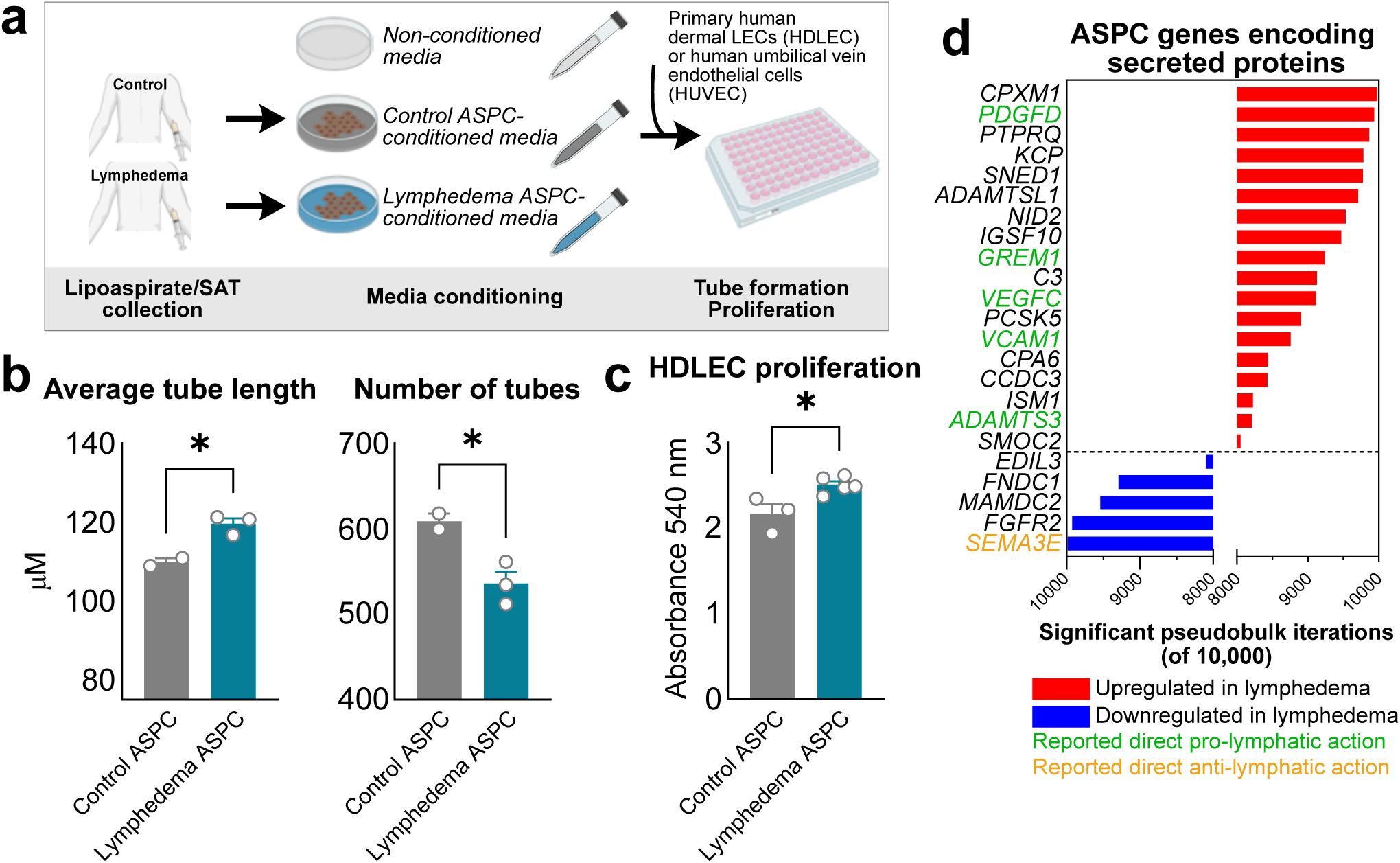
Lymphedema ASPC-secreted factors promote lymphatic endothelial tube elongation and proliferation *ex vivo*. *a*, Experimental paradigm for conditioned media experiments; *b*, HDLEC tube length and number at 18 hours of treatment with conditioned media. Each point represents conditioned media derived from an individual subject, and is the mean of 5 technical replicates; *c*, HDLEC BrdU ELISA absorbance after 24 hours of treatment with conditioned media. Each point represents conditioned media derived from an individual subject, and is the mean of 4 technical replicates; *d*, ASPC pseudobulk up- and down-regulated genes encoding secreted proteins; Statistical analyses performed using independent samples t-test; ns: not significant; *, p<0.05

HDLECs express *VEGFR3*, the primary receptor for VEGF-C, as compared to blood endothelial cells such as HUVECs, which express higher levels of *VEGFR2* (**Extended Data Figure 7c**). Although VEGF-C can also induce signaling via VEGFR-2^71^, when fully processed by CCBE1 the effect of VEGF-C on LECs is significantly more potent than on blood angiogenesis^72^. Treatment of HDLECs with recombinant human VEGF-C results in fewer, longer tubes in a tube formation assay (**Extended Data Figure 7d**), consistent with the lymphatic response to lymphedema ASPC-conditioned media treatment.

Based on ASPC pseudobulk, a number of secreted pro-lymphangiogenic candidates are upregulated in lymphedema (**Figure 8d**), including *PDGFD*, *VEGFC* and *ADAMTS3* as previously discussed, as well as *GREM1*^73^ and *VCAM1*^74^, which have previously reported pro-lymphatic activities. Furthermore, ASPCs downregulate *SEMA3E*, which serves as a repellent against LEC migration^75^. It is possible that the VEGF-C in lymphedema is more highly-processed, and therefore more active toward VEGFR3^71^, given co-expression of processing genes *CCBE1* and *ADAMTS3*. These data indicate that ASPCs upregulate the expression of pro-lymphatic secreted proteins to directly augment lymphangiogenesis in response to lymphatic insufficiency.

## Discussion

Lymphedema is a complex, highly prevalent disease with a poorly-understood pathophysiology defined by both lymphatic dysfunction and dysregulated adipocyte differentiation. Due to challenges in modeling chronic lymphedema in animals, detailed characterization of affected human tissue is essential to developing insights into the underlying mechanisms of disease. In this study, we performed single-nucleus RNA sequencing on lymphedema-associated SAT and identified disease-specific cell states that are associated with adipocyte differentiation and lymphangiogenesis.

In lymphedema, a population of adipocytes expressing *SAA1* and *SAA2* becomes dramatically expanded, a phenomenon also observed in obesity-related adipose deposition^24,76^. However, while SAA expression is correlated with larger adipocyte size in obesity, we found lymphedema-associated adipocytes to be smaller than in control tissue, indicating that adipocyte hyperplasia likely also contributes to adipose expansion in lymphedema. To compensate for chronically disinhibited differentiation, the most stem-like ASPCs, including the DPP4^+^ ASPC^CD55^ population, upregulate genes associated with proliferation and Wnt signaling, which may serve to maintain the adipocyte precursor pool, though we were not able to test *in vivo* ASPC proliferation in this model. Our models of intracellular interactions and downstream signaling suggest that ASPCs may be targeted by IGF-2 and BMP-2 to potentiate these effects.

We also characterized the expansion of the lymphatic vasculature, particularly capillaries, in lymphedema. While bulk RNA sequencing of whole adipose tissue has demonstrated upregulated expression of *PDPN*, *FLT4* (VEGFR-3), and *VEGFC* in lymphedema^31,77,78^, suggesting a pro-lymphangiogenic environment, the source of the factors regulating this process had not yet been identified. In this study, we have defined a previously undescribed lymphedema-associated *GRIA1*^+^ ASPC population that expresses the master lymphangiogenic factor *VEGFC*, its processing machinery, and other secreted agents with known pro-lymphatic actions. Culture media conditioned with both unsorted lymphedema ASPCs and GRIA1+ sorted ASPCs contained more mature VEGF-C and promoted LEC tube elongation and proliferation in an *in vitro* model, indicating that lymphedema-associated ASPCs are programmed to promote lymphangiogenesis. We also noted increased blood vessel density and expression by Ad^SAA^ of *HIF1A* and *VEGFA*. Given the tight bi-directional relationship between angiogenesis and adipose tissue development^79^, the robust vascularization of lymphedema SAT should be further explored for a potential role in directly regulating adipocyte differentiation.

It is difficult to identify a single regulator of ASPC^GRIA1^ activity, as our analysis indicates the role of multiple pathways. We found that TGFβ signaling genes are induced in lymphedema ASPC populations, and TGFβ-1 treatment is known to stimulate *VEGFC* expression in multiple cell types^80,81^. Wnt signaling is also upregulated in ASPC^GRIA1^; Wnt signaling was required for *vegfc* expression in a zebrafish model of lymphangiogenesis^82^, and a study of human medulloblastoma found higher *VEGFC* expression in tumors with aberrant activation of the Wnt pathway compared to other subtypes^83^. We also noted an increase in predicted IGF signaling; IGF1R blockade has recently been reported to reduce VEGF-C production and improve secondary lymphedema in a mouse model^44^, while both IGF-1^84,85^ and IGF-2^86^ are critical for adipocyte progenitor proliferation and differentiation, suggesting this axis may impact both lymphatic function and adipogenesis. Exposure to VEGF-C itself has also been demonstrated to induce ASPC *VEGFC* expression and release^87^, and may promote additional pro-lymphangiogenic mechanisms, such as secretion of exosomal microRNAs ^88^.

There has been interest in using ASPCs therapeutically for the treatment of secondary lymphedema^89^, and the capacity of ASPCs to secrete lymphangiogenic factors, including VEGF-C, has been described in non-pathogenic settings^90^. Small trials of autologous ASPC transfer in humans with lymphedema, however, have thus far failed to show benefit with respect to limb volume or lymphoscintigraphic measures^91,92^. In our study population, lymphedema limb volume by perometry was reduced with increased prevalence of ASPC^GRIA1^, but not other ASPC subpopulations, suggesting that the potential effectiveness of ASPCs to mitigate lymphedema, particularly the fluid component, may rely on being activated toward a pro-lymphangiogenic state. Further characterization of the upstream signaling that regulates this specialized ASPC identity should be the focus of future investigation.

In summary, this atlas provides a valuable resource to thoroughly interrogate the cell populations and activities that define secondary lymphedema in order to design mechanistic studies with the ultimate goal of developing clinically useful therapies and diagnostic biomarkers. Limitations of this study include limited sample size and lack of power to directly compare lymphedema etiologies, effect of subject sex, or limb affected. Subjects were recruited from a single center, and all samples used for sNuc-seq were obtained from Caucasian, non-Hispanic individuals.

## Online methods

### Subject enrollment and adipose tissue sample collection

All adult subjects with unilateral, fat-predominant (stage 2 or higher) lymphedema undergoing surgical debulking at BIDMC with a BMI between 18 and 35 kg/m^2^ were screened for inclusion in this study, and were excluded for presence of the following: history of diabetes mellitus or use of glucose-lowering or weight loss medications; use of anti-estrogen agents, glucocorticoids, immunosuppression, or chemotherapy within the past 6 months; history of HIV/AIDS; or active malignancy. Subjects provided written informed consent preoperatively and were not compensated for their participation (BIDMC IRB 2021P000364). Subcutaneous adipose tissue for sequencing, approximately 50 mg, was resected en bloc at the start of the surgical case under general anesthesia via a 4 mm incision and snap frozen immediately in liquid nitrogen. Biopsies for histological slide preparation were obtained using a 6 mm biopsy punch (Fisher Scientific) and placed on ice. Lipoaspirate, approximately 20 mL, was obtained as per standard of care at the beginning of the case using power assisted liposuction as described elsewhere^93^.

### Single-nucleus RNA sequencing and data analysis

SAT sNuc-seq was performed as previously described^94^. Briefly, adipose samples were homogenized in TST buffer using a gentleMACS Dissociator (Miltenyi Biotec). Lysate was filtered through 40 µm and 20 µm nylon filters (CellTreat) and washed. After the second wash, each sample was incubated with NucBlue (Thermofisher Scientific) as well as an individual hashtag antibody (BioLegend) for 45 minutes. After incubation, nuclei were washed and flow sorted together into RT Reagent B (10x Genomics) using a Beckman Coulter MoFlo Astrios EQ with a 70 μm nozzle. After sorting, samples were immediately loaded on the 10x Chromium controller (10x Genomics) according to the manufacturer’s protocol. Single Cell 3’ v3.1 chemistry was used to process all samples and cDNA and gene expression libraries were generated according to the manufacturer’s instructions (10x Genomics). Gene expression libraries were multiplexed and sequenced on the Nextseq 500 (Illumina).

Raw sequencing reads were processed into digital expression matrices (DGE) using CellRanger (10x Genomics, version 6.1.2) and using annotation GENCODE-v44. DemuxEM32 (version 0.1.6) was used to sort cells into individual samples based on their antibody hashtags. Ambient RNA was removed from the processed DGEs using CellBender (version 0.2.0), and doublet scores were calculated using both scDblFinder (version 1.2.0) and scds (version 1.4.0). Cells were removed as doublets if they were both determined to be a doublet using scDblFinder and if they had a scds hybrid score > 1.5. Cells with < 800 UMIs were removed from the dataset, as were cells with > 10% of mitochondrial reads. Genes found in fewer than 2 cells were removed. Seurat (version 4.1.1) was used as the single-cell clustering and analysis framework. The raw count data was normalized (scaled and centered) per-sample using the function “SCTransform”, using mitochondrial read content (percentage), ribosomal read content, and the *S* and *G2* phase scores (per the Seurat function “CellCycleScoring”) as regression variables. Next, all sample-wise objects were integrated using CCA.. To subcluster, objects were subset by general cell type and re-analyzed; sub-objects were split by sample, rescaled and then reintegrated, and the clusters were recalculated. After the initial round of subclustering, subclusters with high doublet scores and/or high percentages of mitochondrial reads were removed and the data was, once again, reintegrated and clusters were recalculated.

Reference mapping was performed between the reported dataset and the human subcutaneous dataset from Emont et. al. (2022) using Seurat multimodal reference mapping. To annotate cell types, calculated clusters were evaluated for similarity to previously annotated clusters. If clusters mapped exactly, they were annotated as the same cell type. If not, they were evaluated for distinct marker genes and annotated as a new cluster. All clusters without distinct names were renamed to be defined by a top marker gene.

To calculate pseudobulk counts, objects for individual cell types were split by sample and randomly subset to contain the same number of cells per sample. Differential expression analysis (DEA) was run on pseudobulk samples using edgeR (version 3.30.3). To reduce the variation introduced by the random subsampling, subsampling and DEA were performed 10,000 times; significantly regulated genes used on plots and for correlation analysis were defined as those with log_2_ fold change greater than 0.5 and FDR less than 0.1 in at least 9,000 iterations. To plot, heatmaps were generated using pseudobulk sums of all cells in each sample, and edgeR was used to calculate cpms from the pseudobulk counts. PCA plots were generated using a randomly subsampled pseudobulk dataset. Heatmaps were plotted using pheatmap (version 1.0.12).

Predicted intercellular interactions and downstream signaling pathways were determined using NicheNet^38^ and CellChat^42^. Signature scores were calculated using Vision 3.0.0^95^ using gene lists from the Molecular Signatures Database^26^ as listed in **Supplementary Table 4**. Comparisons between groups were performed using a Wilcoxon rank sum test.

### MRI imaging analysis

ImageJ was used to manually segment and threshold fat on fat-weighted MR sequences of the bilateral extremities at the approximate level of the biopsies in order to calculate fat volume, fluid volume, fat fraction and total subcutaneous area in the affected and unaffected contralateral extremity.

### Histology and immunofluorescent labeling

Tissue biopsies were immediately placed on ice, washed with PBS, and fixed in 10% formalin overnight at 4°C, followed by paraffin embedding, sectioning, and staining with hematoxylin and eosin (H&E). Unstained sections for immunofluorescence were deparaffinized and rehydrated by heating to 55°C and sequential washing in xylene, 100%, 95%, 70%, 50% ethanol and deionized water. Antigen retrieval was performed in sub-boiling 10 mM sodium citrate buffer (pH 6.0) for 10 min. Sections were washed with 1% donkey serum in PBS with 0.4% Triton X-100 (PBS-T) twice and blocked with 5% donkey serum in PBS-T for 30 minutes. Sections were then incubated with the primary antibodies diluted in 1% donkey serum PBS-T at room temperature for 1 h and then at 4°C overnight. Primary antibody and dilutions used for labeling were: Alexa Fluor 488 rabbit anti-PLIN1 (1:100, Cell Signaling, 29138S), goat anti-LYVE 1 (1:100, R&D Systems, AF2089), and rabbit anti-CD31 (1:200, Proteintech, 11265-1-AP). Samples were subsequently incubated with secondary antibodies diluted in PBS-T at room temperature for 1 hour. Secondary antibodies used were: Alexa Fluor 647 donkey anti-goat (1:250, Thermo Scientific, A-31571), Alexa Fluor 568 donkey anti-mouse (1:250, Thermo Scientific, A-19937), Alexa Fluor 568 donkey anti-rabbit (1:250, Thermo Scientific, A-10042) and Hoechst 33342 (1:1000, Invitrogen, H3570). Images were obtained using a fluorescent slide scanner and Zeiss LSM 880 laser scanning confocal system running Zen Black 2.3 software.

### Adipocyte size, lymphatic vessel quantification, and vascular density measurement

To measure adipocyte size, H&E and confocal BODIPY-labeled images were analyzed using Adiposoft v1.16^96^ in ImageJ Fiji. All counts were manually verified, and adipocytes inaccurately delineated in automatic mode were manually adjusted. Lymphatic vessels were measured using ImageJ Fiji software. To quantify lymphatic vessels, tissue sections were fluorescently immunolabeled for CD31 and LYVE1 as above. Two to three equally-powered fields per sample were evaluated for quantification. Lymphatic vessels were identified as vascular structures double-labeled for LYVE1 and CD31, and these were manually measured length-wise using ImageJ, or for vessels cut in cross-section, measurement was performed along the longest axis. To measure overall vascular density, tissue sections immunolabeled for CD31+ were analyzed using Vessel Analysis plugin^97^ for ImageJ and vascular length density measurements were obtained.

### Cell culture

Fresh lipoaspirate was incubated with collagenase type II (1 mg/ml) (Sigma-Aldrich) in digestion buffer (Hanks’ balanced salt solution with calcium and magnesium supplemented with 0.5% bovine serum albumin) on a shaker water bath incubator at 37°C for 20 minutes with vigorous shaking by hand every 5 minutes. EDTA, pH 8.0 at a concentration of 10 mM was then added for an additional 5 minutes, after which wash media (DMEM+glutamax with 10% fetal bovine serum (FBS) and 1% penicillin-streptomycin) was added. Cells were filtered through a 100 μm filter and washed, and if flow cytometry was to be performed, incubated with ACK lysis buffer for 5 minutes. SVF was then washed and filtered through 40 μm filter followed by plating on multiwell plates in growth media (DMEM+glutamax, 15% FBS, 1% penicillin-streptomycin). Human primary dermal lymphatic endothelial cells (HDLEC, Cell Biologics, H-6064) and human umbilical vein endothelial cells (HUVEC, gift from Shingo Kajimura) were maintained on gelatin-coated flasks in complete human endothelial cell media (Cell Biologics, H-1168).

### RT-quantitative PCR

Cells were collected in Trizol as described above. Total RNA was isolated using the E.Z.N.A. Total RNA Kit II (Omega Bio-Tek) per manufacturer’s instructions. RNA concentration was quantified using NanoDrop One (Thermo Scientific) and up to 1 µg of RNA was reverse-transcribed using High-Capacity cDNA Reverse Transcription Kit (Thermo Fisher). SYBR Green PCR Master Mix (Thermo Fisher) and gene primers (**Supplementary Table 54**) were used to perform RT-qPCR using QuantStudio 6 Flex Real-time PCR system. Expression was normalized to the housekeeping gene *ACTB*. Results are presented as fold change (2^−ΔCT^) and comparisons between HDLEC and HUVEC were performed with two-tailed Student’s T-test.

### Conditioned media (CM) generation, and *in vitro* tube formation and proliferation assays

ASPCs at passages 3-5 were grown to confluence in 6 well plates. DMEM+glutamax supplemented with 0.5% FBS and 1% pen-strep, 2 mL per well, was conditioned at 37°C for 48 hours, centrifuged at 10,000 G for 10 minutes at 4°C to remove cellular debris, and stored at −80°C.

To perform tube formation assays, growth-factor reduced Matrigel (Corning) was added to 96 well plates, followed by addition of 1.5 x 10^4^ HDLECs or HUVECs per well in complete EC media. Imaging was performed at 18 hours after plating, and tube networks were analyzed using Angiogenesis Analyzer for ImageJ^98^. For CM experiments, CM was mixed 1:1 with complete EC media at the time of cell plating. Recombinant human VEGF-C (BioLegend, 589702) was used at a concentration of 50 ng/mL.

To perform proliferation assay, 5 x 10^3^ HDLECs per well were plated on gelatin coated 96-well plates. After 2 hours, cells were washed with PBS twice and media was switched to serum-free basal endothelial cell media (Cell Biologics, H1168b) for 18 hours. Following serum-starvation, media was switched to CM mixed 1:1 with complete EC media with BrdU label for 24 hours and assayed using BrdU Cell Proliferation Assay (Sigma-Aldrich, QIA58).

### Statistics

Statistics were performed using GraphPad Prism 10. Population proportion scores and MRI feature comparisons between lymphedema and control samples were performed as paired two-sided Students t-tests. In conditioned media experiments, to compare control- and lymphedema-conditioned samples, we first calculated the means for each biological replicate (by averaging over technical replicates). We then implemented an independent samples t-test to test the difference in the presented measure between these two groups. Error bars represent the standard error of the mean.

## Data availability

Raw and processed sNuc-seq data will be available on GEO at the time of publication. Processed data are available on figshare (reviewer link: https://figshare.com/s/9ee8d48a45647fe1e6f5). Data from the Emont et. al. 2022 study is publicly available: GSE176067.

## Acknowledgements

We gratefully acknowledge the BIDMC Department of Surgery FIRST program for their assistance developing and maintaining the IRB protocol for subject enrollment and sample collection, and the Boston Area Diabetes Endocrinology Research Center Functional Genomics and Bioinformatics Core (supported by NIH P30 DK135043) for performing sNuc-seq. GPW is supported by NIH K08 DK132413, EDR is supported by NIH R01 DK126789 and RC2 DK116691, MPE is supported by NIH K01 DK134806, and TPP is supported by NIH R01 CA284372 and DoD HT94252410100.

---

**Extended Data Figure 1.**
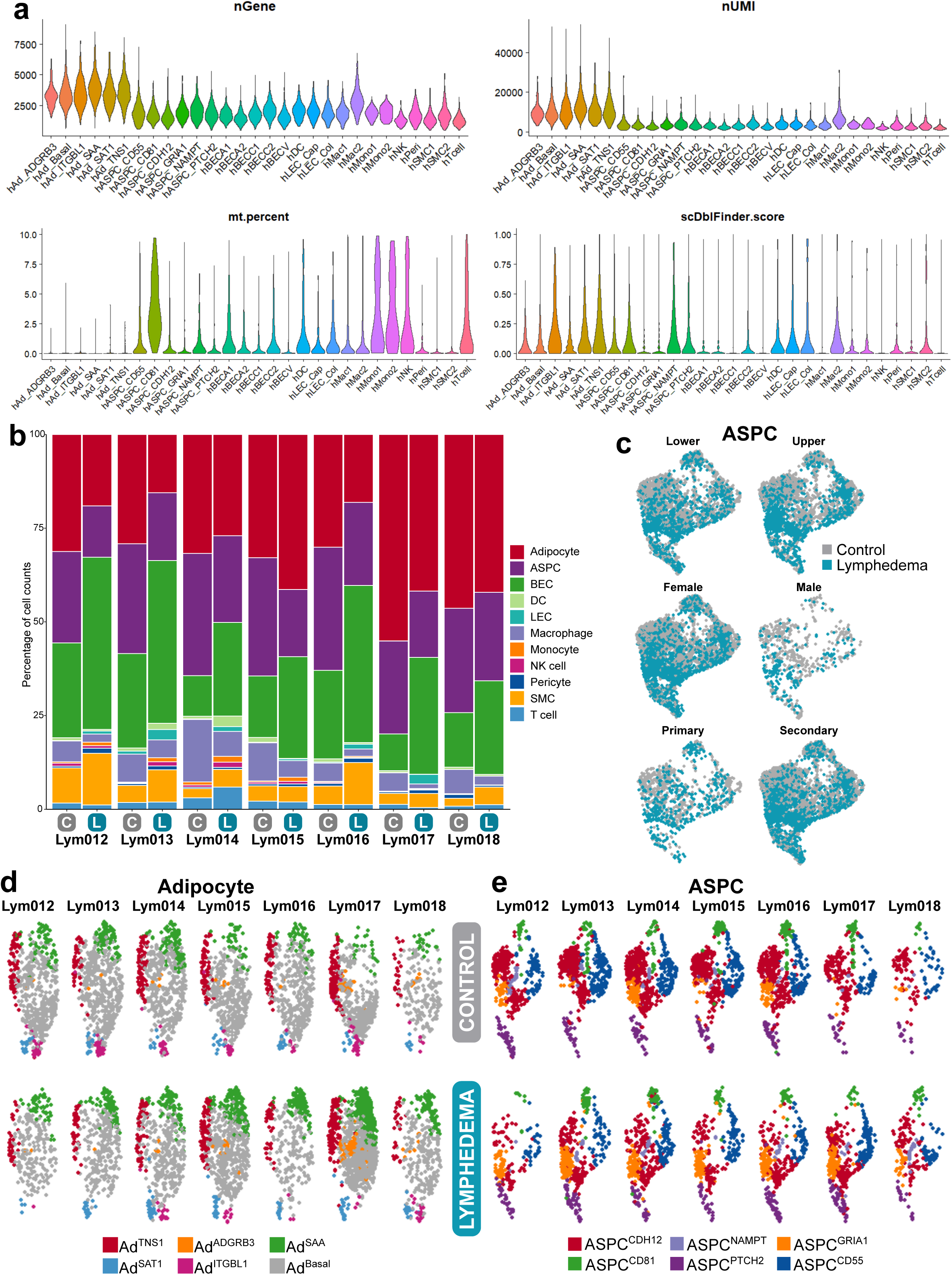
*a,* Quality control metrics for sNuc-seq data by subpopulation, including gene and UMI counts, mitochondrial transcript content, and Scrublet score for doublets; *b*, Cell population proportions split by sample; *c*, ASPCs split by biological variable and grouped by disease state; *d-e*, UMAP plots of adipocytes and ASPCs split by sample. C, control; L, lymphedema.

**Extended Data Figure 2.**
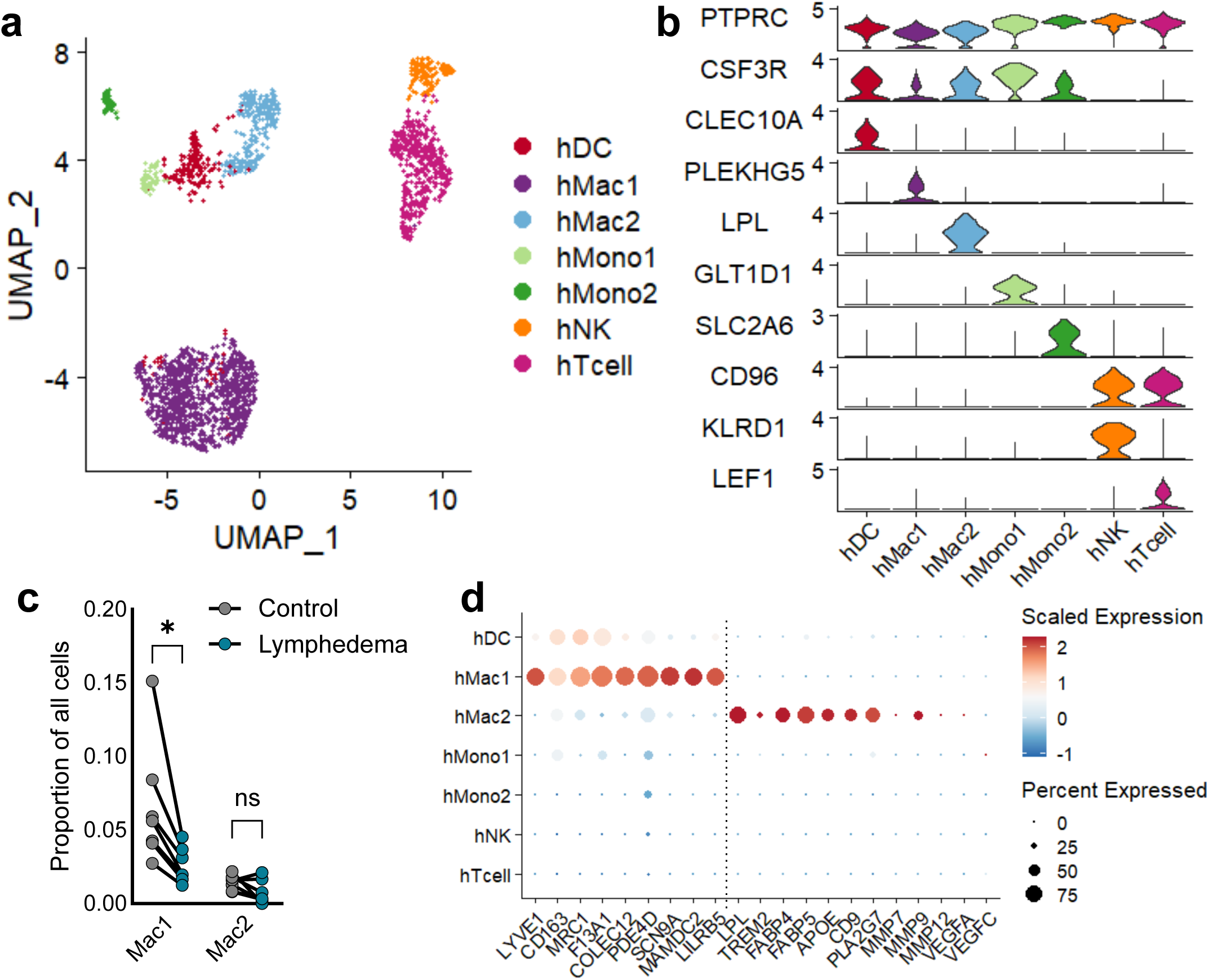
*a,* UMAP plot of immune cell subpopulations; *b*, Log-normalized expression of unique marker genes by subpopulation; *c*, Macrophage subpopulations as a proportion of all cells; *d*, Dot plot of expression of genes relevant to macrophage subpopulation identity and function. Statistical analysis performed using two-tailed paired Student’s t-test; *, p<0.05.

**Extended Data Figure 3.**
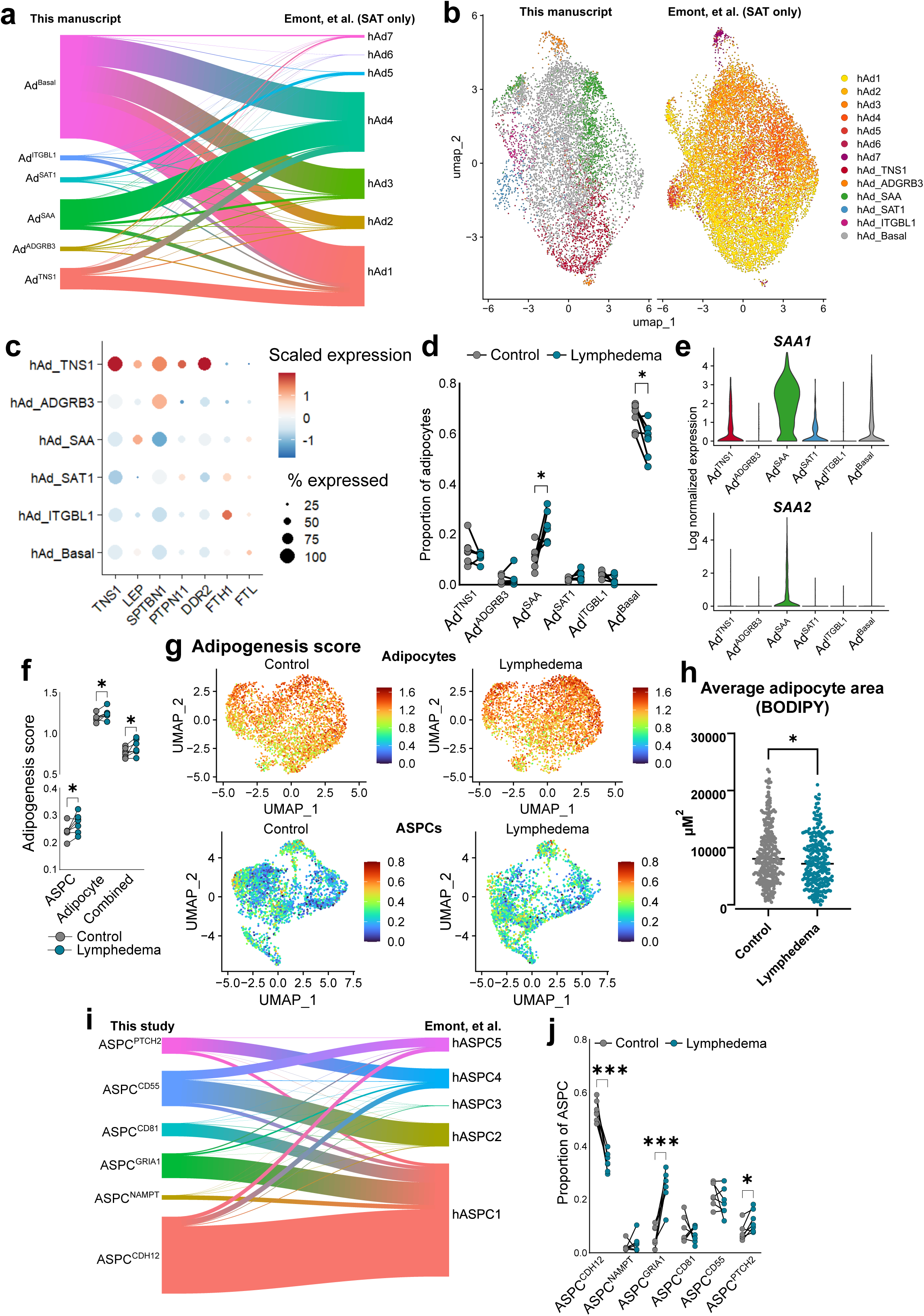
*a,* Adipocyte label transfer and *b*, integration with Emont, et. al. reference subcutaneous adipocyte atlas; *c*, Expression of selected Adipo^Lep^ marker genes described by Bäckdahl, et al. *d*, Adipocyte subpopulation proportions; *e*, *SAA1* and *SAA2* log-normalized expression across adipocyte subpopulations; *f*, Adipogenesis score disaggregated by individual and cell type; *e*, Quantification of adipocyte size on BODIPY-labeled SAT whole mount; *f,* ASPC label transfer to Emont, et. al. reference populations; *g*, ASPC subpopulation proportions. Statistical analysis performed using paired (f, j) or unpaired (h) two-tailed Student’s t-tests; *, p<0.05; ***, p<0.001. SAT, subcutaneous adipose tissue

**Extended Data Figure 4.**
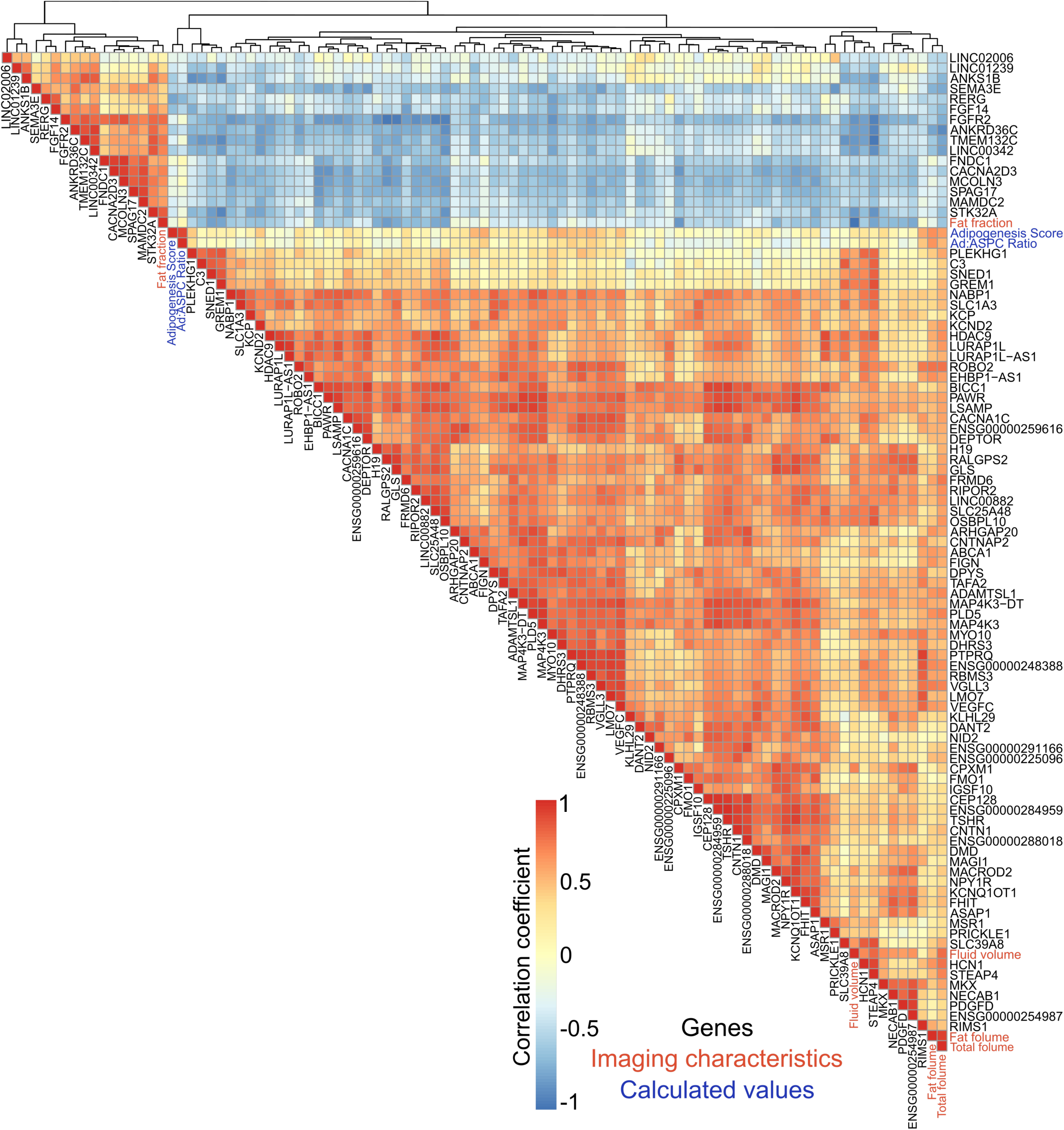
Heatmap of Pearson correlations between genes identified in ASPC pseudobulk as well as with radiographically determined imaging characteristics (in red) and calculated adipogenesis score and Ad:ASPC ratio (as an additional surrogate of adipocyte differentiation, in blue).

**Extended Data Figure 5.**
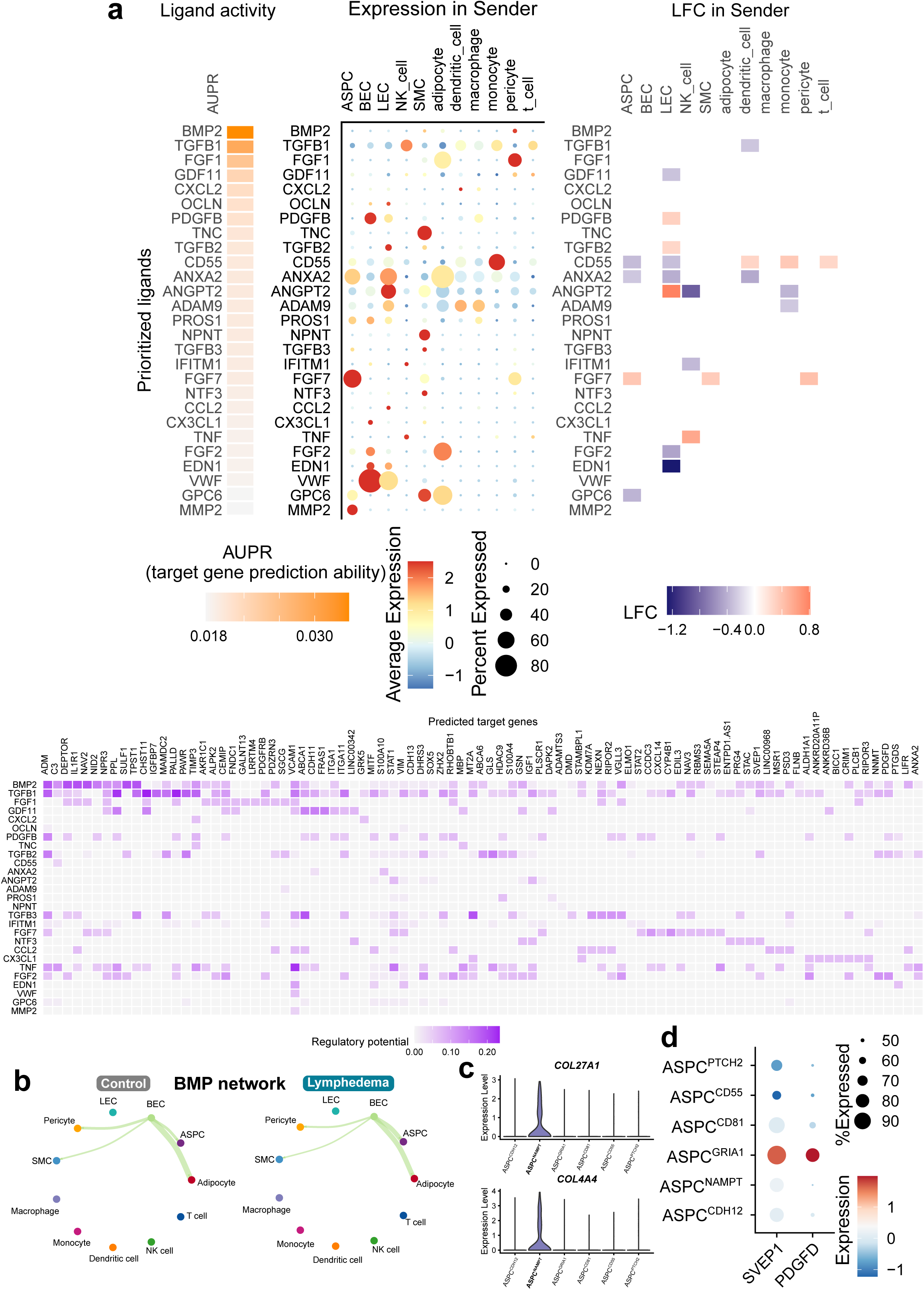
*a,* NicheNet analysis (sender-focused) of ASPC ligand-receptor interactions; *b,* CellChat intercellular interaction modeling of the BMP signaling network between cell types; *c*, Selected collagen gene expression of ASPC^NAMPT^; *d*, Expression of additional selected pro-lymphatic secreted proteins by ASPC subpopulation. AUPR, area under the precision-recall curve; LFC, log-fold change.

**Extended Data Figure 6.**
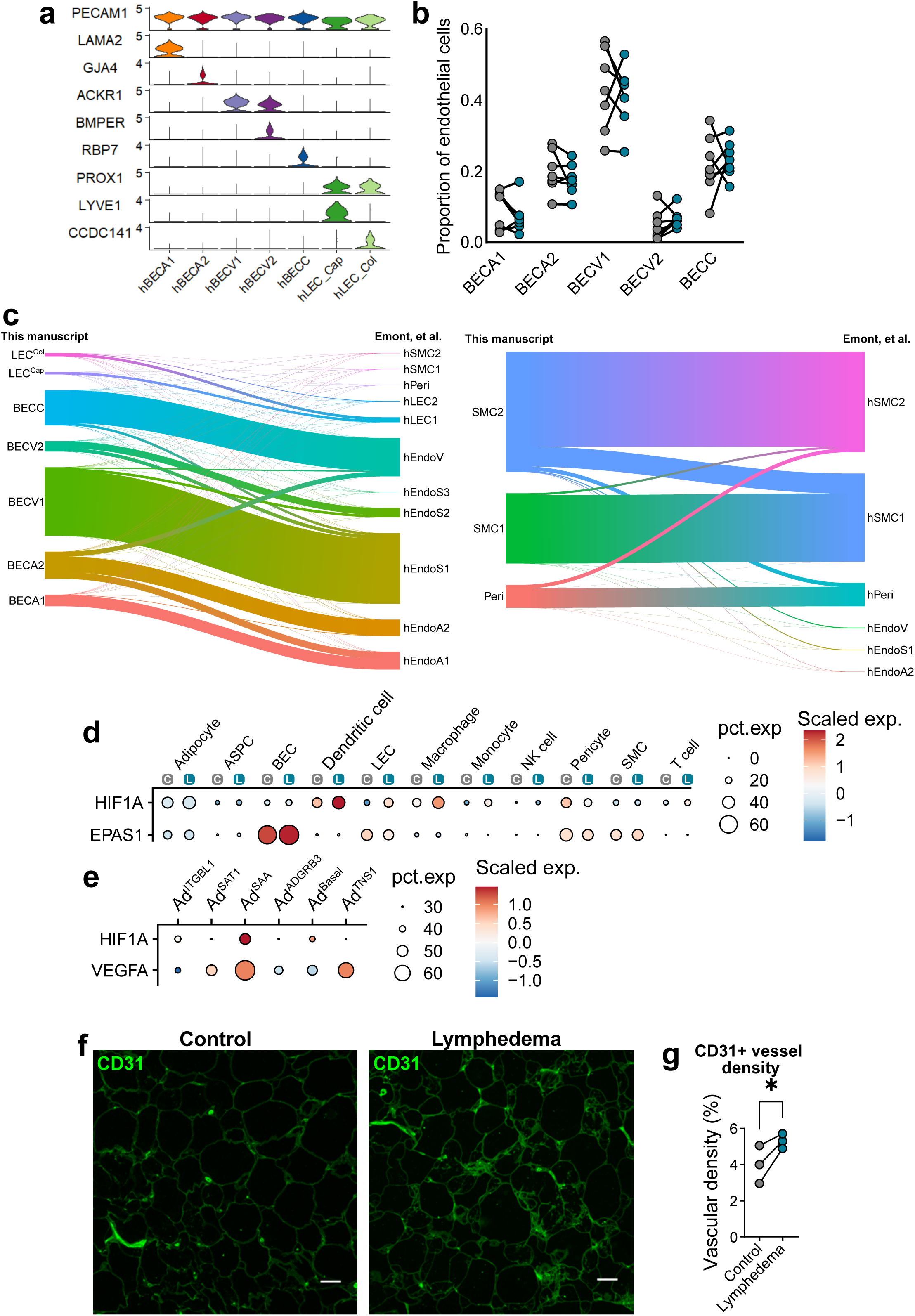
*a*, Selected marker genes for endothelial cell populations; *b*, BEC subpopulations as a proportion of total endothelial cells; *c*, Endothelial cell, SMC, and pericyte label transfer to Emont, et. al. reference populations; *d,* Expression of hypoxia-inducible factors across all cell populations; *e*, *HIF1A* and *VEGFA* expression within adipocyte subpopulations; *f*, Representative immunofluorescence for CD31 of paired SAT biopsies; *g*, Vascular density quantification. C, control; L, lymphedema. Statistical analysis performed using paired two-tailed Student’s t-test; *, p<0.05.

**Extended Data Figure 7.**
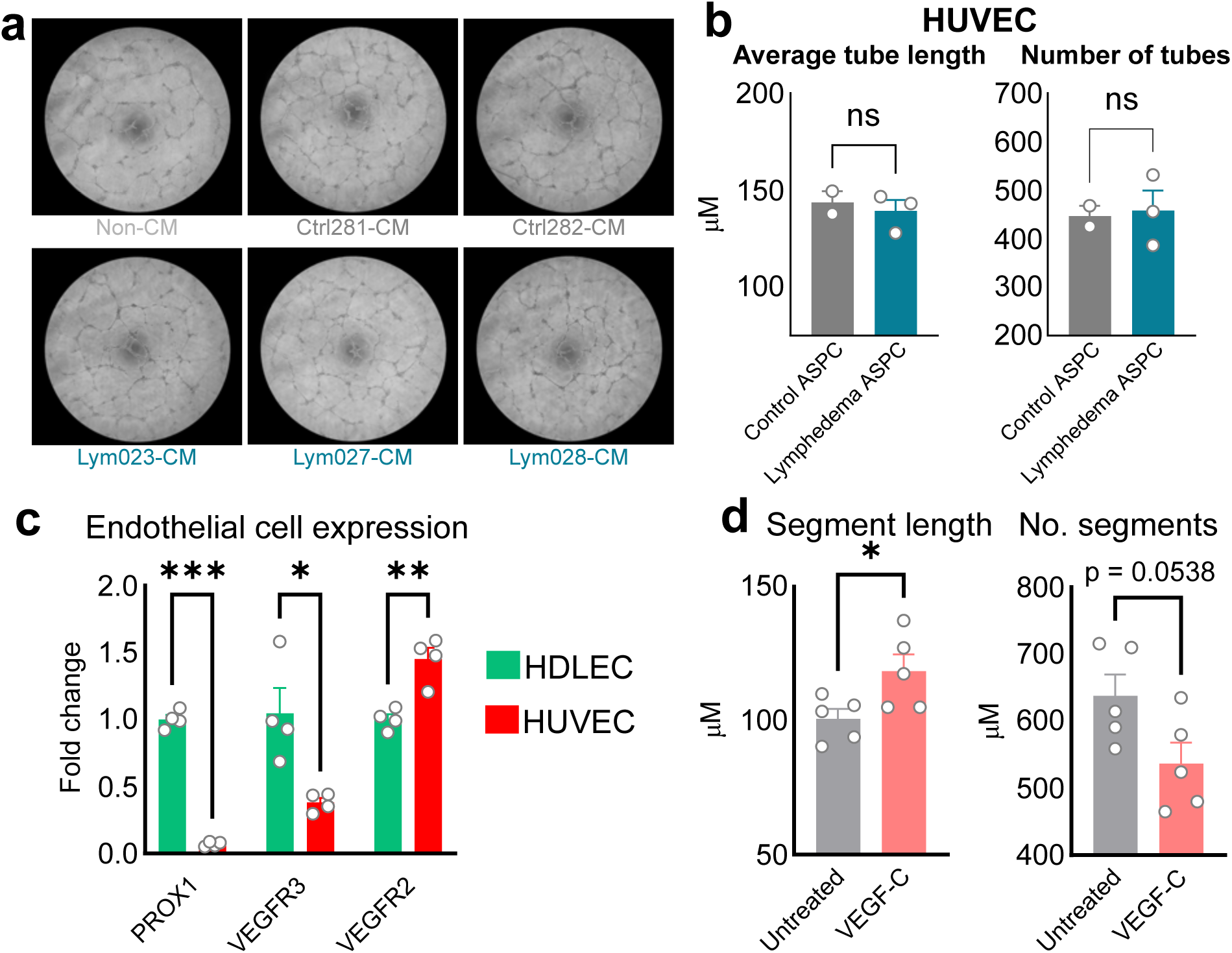
*a,* Representative light microscopy of Matrigel tube formation assays at 18 hours; *b*, HUVEC tube parameters at 18 hours of treatment. Each point represents conditioned media derived from an individual subject, and is the mean of 5 technical replicates; *c*, qPCR for selected endothelial identity genes; *d*, HDLEC tube parameters at 18 hours of treatment with recombinant human VEGF-C; Statistical analyses performed using unpaired two-tailed Student’s t-test; *, p<0.05; **, p<0.01; ***, p<0.001.

## Notes

### Competing Interest Statement

The authors have declared no competing interest.

### Summary of Updates

Ad2 and Ad3 subpopulations combined (Figure 2a-c); Figure 2n-m removed; Figure 4h-i removed; New Figure 5d; Figure 6 (new Figure 8) updated analysis in panel 6c (now 8b) and added analysis in panel 8c, removed panels 6d-g and added new panel 8d; Extended Data Figure 1 added with QC metrics; Extended Data Figure 6g analysis added; Extended Data Figure 5 panel e removed; updated supplemental files; other formatting and organizational changes to improve readability.

