## Supplemental Tables for "Single-nuclear transcriptomics of lymphedema-associated adipose reveals a pro-lymphangiogenic stromal cell population"

**Supplementary Table 1: Subject characteristics**

| **Subject ID** | **Age** | **Sex** | **BMI** | **Lymphedematous limb** | **Lymphedema etiology** | **Application** |
| --- | --- | --- | --- | --- | --- | --- |
| Lym012 | 63 | M | 23.8 | Lower | Secondary | sNuc-seq, MRI, and perometry |
| Lym013 | 64 | F | 25.1 | Upper | Secondary | sNuc-seq, MRI, and perometry |
| Lym014 | 55 | F | 24.1 | Lower | Idiopathic/primary | sNuc-seq, MRI, and perometry |
| Lym015 | 77 | F | 33.1 | Upper | Secondary | sNuc-seq, MRI, and perometry |
| Lym016 | 58 | F | 31.9 | Lower | Secondary | sNuc-seq, MRI, and perometry |
| Lym017 | 75 | F | 22.4 | Upper | Secondary | sNuc-seq, MRI, and perometry |
| Lym018 | 54 | F | 19.1 | Lower | Secondary | sNuc-seq, MRI, and perometry |
| Lym029 | 59 | F | 29.3 | Upper | Secondary | Histology and immunofluorescence |
| Lym030 | 58 | F | 26 | Upper | Secondary | Histology and immunofluorescence |
| Lym031 | 57 | F | 26.6 | Lower | Idiopathic/primary | Histology and immunofluorescence |
| Lym023 | 71 | F | 33.5 | Upper | Secondary | Conditioned media (tube formation) |
| Lym027 | 79 | F | 30.7 | Upper | Secondary | Conditioned media (tube formation and proliferation) |
| Lym028 | 70 | F | 29.9 | Lower | Idiopathic/primary | Conditioned media (tube formation and proliferation) |
| Lym032 | 65 | F | 25.1 | Upper | Secondary | Conditioned media (proliferation) |
| Lym033 | 34 | F | 22.3 | Lower | Idiopathic/primary | Conditioned media (proliferation) |
| Lym034 | 71 | F | 27.2 | Lower | Idiopathic/primary | BODIPY adipocyte quantification and conditioned media (proliferation) |
| Ctrl281 | 68 | F | 29.0 | n/a. Control sample lower extremity SAT | n/a | Conditioned media (tube formation and proliferation) |
| Ctrl282 | 48 | F | 31.0 | n/a. Control sample abdominal SAT | n/a | Conditioned media (tube formation and proliferation) |
| Ctrl283 | 37 | F | 25.1 | n/a. Control sample abdominal SAT | n/a | Conditioned media (proliferation) |

**Supplementary Table 2: MRI and perometry measurements**

| **Subject ID** | **Control fat area (mm^2^)** | **Control fluid area (mm^2^)** | **Control total area (mm^2^)** | **Lymphedema fat area (mm^2^)** | **Lymphedema fluid area (mm^2^)** | **Lymphedema total area (mm^2^)** | **Perometry (% volume difference between limbs)** |
| --- | --- | --- | --- | --- | --- | --- | --- |
| Lym012 | 2207.48 | 407.597 | 2615.077 | 3871.767 | 1014.553 | 4886.32 | 40.4 |
| Lym013 | 2343.291 | 197.799 | 2541.09 | 2690.776 | 889.203 | 3579.979 | 22 |
| Lym014 | 1800 | 245.215 | 2045.215 | 3131.543 | 2342.285 | 5473.828 | 66 |
| Lym015 | 2690.776 | 89.062 | 2779.838 | 6190.625 | 1426.562 | 7617.187 | 36 |
| Lym016 | 3184.277 | 373.535 | 3557.812 | 4100.977 | 1947.656 | 6048.633 | 38 |
| Lym017 | 3079.688 | 200 | 3279.688 | 5596.875 | 970.312 | 6567.187 | 40 |
| Lym018 | 2296.209 | 513.305 | 2809.514 | 3754.143 | 1884.826 | 5638.969 | 61 |

**Supplementary Table 3: Cell type proportions by individual**

|  |  | **Adipocyte** | **ASPC** | **BEC** | **SMC** | **Macroph.** | **T-cells** | **DCs** | **LEC** | **Pericytes** | **Mono.** | **NK cells** |
| --- | --- | --- | --- | --- | --- | --- | --- | --- | --- | --- | --- | --- |
| **Lym012** | Control | 0.3122 | 0.244006 | 0.253063 | 0.093767 | 0.055408 | 0.016516 | 0.007459 | 0.001066 | 0.004795 | 0.003729 | 0.007991 |
|  | Lymph. | 0.19058 | 0.136957 | 0.45942 | 0.137681 | 0.021739 | 0.011594 | 0.004348 | 0.007971 | 0.013768 | 0.00942 | 0.006522 |
| **Lym013** | Control | 0.291667 | 0.292996 | 0.251773 | 0.045213 | 0.074911 | 0.018174 | 0.008422 | 0.007979 | 0.004876 | 0.001773 | 0.002216 |
|  | Lymph. | 0.155366 | 0.180938 | 0.434328 | 0.086013 | 0.047656 | 0.019372 | 0.01666 | 0.027509 | 0.010074 | 0.010849 | 0.011236 |
| **Lym014** | Control | 0.317426 | 0.326005 | 0.108311 | 0.024665 | 0.166756 | 0.030563 | 0.006971 | 0.001609 | 0.003753 | 0.008043 | 0.005898 |
|  | Lymph. | 0.270227 | 0.231392 | 0.24973 | 0.046926 | 0.066343 | 0.059331 | 0.028047 | 0.012945 | 0.005394 | 0.015642 | 0.014024 |
| **Lym015** | Control | 0.329004 | 0.315399 | 0.163884 | 0.040816 | 0.102041 | 0.021645 | 0.012987 | 0.001237 | 0.004947 | 0.003092 | 0.004947 |
|  | Lymph. | 0.413495 | 0.179354 | 0.271626 | 0.040946 | 0.044406 | 0.019608 | 0.001153 | 0.004614 | 0.00692 | 0.010957 | 0.00692 |
| **Lym016** | Control | 0.300798 | 0.328422 | 0.236341 | 0.04911 | 0.04911 | 0.012891 | 0.008594 | 0.002455 | 0.006753 | 0.002455 | 0.003069 |
|  | Lymph. | 0.181267 | 0.221488 | 0.419284 | 0.111846 | 0.019284 | 0.012672 | 0.004408 | 0.012672 | 0.01157 | 0.003306 | 0.002204 |
| **Lym017** | Control | 0.550943 | 0.248113 | 0.098113 | 0.029245 | 0.049057 | 0.013208 | 0.004717 | 0.000943 | 0.003774 | 0.001887 | 0 |
|  | Lymph. | 0.417916 | 0.176965 | 0.311517 | 0.037294 | 0.012431 | 0.004753 | 0.000731 | 0.025229 | 0.009506 | 0.001828 | 0.001828 |
| **Lym018** | Control | 0.463617 | 0.278586 | 0.14553 | 0.02079 | 0.064449 | 0.008316 | 0.006237 | 0 | 0.010395 | 0.002079 | 0 |
|  | Lymph. | 0.421053 | 0.236282 | 0.24972 | 0.047032 | 0.023516 | 0.012318 | 0.004479 | 0 | 0.004479 | 0.00112 | 0 |

**Supplementary Table 4: Gene signatures**

| **Signature** | **mSigDB gene set** |
| --- | --- |
| Adipogenesis | Hallmark Adipogenesis |
| Angiogenesis | Hallmark Angiogenesis |
| Lymphangiogenesis | PID Lymphangiogenesis |
| Stem cell proliferation | GOBP Stem cell proliferation |
| IGF | GOBP Insulin-like growth factor receptor signaling pathway |
| Insulin | KEGG Insulin signaling pathway |
| BMP | GOBP Response to BMP |
| TGF-β | Hallmark TGF beta signaling |
| WNT | GOBP Canonical WNT signaling pathway |
| FOXO | PID FOXO pathway |
| Notch | Hallmark Notch signaling |
| Angiopoietin | PID angiopoietin receptor pathway |

**Supplementary Table 5: qPCR primers**

| **Gene** | **Forward primer** | **Reverse primer** |
| --- | --- | --- |
| *ACTB* | GCACAGAGCCTCGCCTT | GTTGTCGACGACGAGCG |
| *FLT4* (VEGFR-3) | AGCCATTCATCAACAAGCCT | GGCAACAGCTGGATGTCATA |
| *KDR* (VEGFR-2) | CCAGCAAAAGCAGGGAGTCTGT | TGTCTGTGTCATCGGAGTGATATCC |
| *PROX1* | CCAGCTCCAATATGCTGAAGACCTA | CATCGTTGATGGCTTGACGTG |
